# Pervasive nuclear envelope ruptures precede ECM signaling and disease onset without activating cGAS-STING in Lamin-cardiomyopathy mice

**DOI:** 10.1101/2023.08.28.555134

**Authors:** Atsuki En, Hanumakumar Bogireddi, Briana Thomas, Alexis Stutzman, Sachie Ikegami, Brigitte LaForest, Omar Almakki, Peter Pytel, Ivan P. Moskowitz, Kohta Ikegami

## Abstract

Nuclear envelope (NE) ruptures are emerging observations in Lamin-related dilated cardiomyopathy, an adult-onset disease caused by loss-of-function mutations in Lamin A/C, a nuclear lamina component. Here, we tested a prevailing hypothesis that NE ruptures trigger pathological cGAS-STING cytosolic DNA-sensing pathway, using a mouse model of Lamin-cardiomyopathy. Reduction of Lamin A/C in cardiomyocytes of adult mice caused pervasive NE ruptures in cardiomyocytes, preceding inflammatory transcription, fibrosis, and fatal dilated cardiomyopathy. NE ruptures were followed by DNA damage accumulation without causing immediate cardiomyocyte death. However, cGAS-STING-dependent inflammatory signaling remained inactive. Deleting *cGas* or *Sting* did not rescue cardiomyopathy. The lack of cGAS-STING activation was likely due to the near absence of cGAS expression in adult cardiomyocytes at baseline. Instead, extracellular matrix (ECM) signaling was activated and predicted to initiate pro-inflammatory communication from Lamin-reduced cardiomyocytes to fibroblasts. Our work nominates ECM signaling, not cGAS-STING, as a potential inflammatory contributor in Lamin-cardiomyopathy.

## INTRODUCTION

Lamin A/C (*LMNA*) are nuclear lamina proteins that provide structural integrity to the nuclear envelope ^1–4^. Mutations in *LMNA* cause a spectrum of degenerative disorders, collectively called laminopathies, including frequent dilated cardiomyopathy (*LMNA*-related DCM) ^5–10^. *LMNA*-related DCM is prevalent adult-onset DCM ^11,12^ that accompanies cardiac conduction disease, fibrosis, heart failure, and mortality ^13–16^. *LMNA*-related DCM is predominantly caused by heterozygous loss-of-function *LMNA* mutations that cause Lamin A/C protein reduction most strongly in cardiomyocytes ^16–20^. Activation of stress response-related signaling, such as MAPK signaling ^21–24^, mTOR signaling ^25^, and DNA damage response signaling ^26,27^, has been reported in *LMNA*-related DCM models. However, the direct pathological alterations caused by Lamin A/C reduction in cardiomyocytes remain undefined.

Reduction of nuclear lamins can cause ruptures of the nuclear envelope (NE ruptures) ^28–31,33–35^. Indeed, NE ruptures have been observed in patients ^17,18,32,36,37^, animal models ^30,31,38–42^, and cell culture models ^30,35^ of *LMNA*-related DCM and laminopathies. However, the extent of NE ruptures in *LMNA*-related DCM and the contribution of this event to the pathogenesis of DCM remain undefined.

A prevailing hypothesis for *LMNA*-related DCM pathogenesis implicates the cGAS-STING pathway as a link between NE rupture, inflammatory signaling, and DCM pathogenesis ^39,40,43–45^. cGAS-STING is a cytosolic DNA-sensing innate immune mechanism, in which cGAS binds to cytosolic DNA and activates cell-autonomous STING-mediated interferon transcription ^46,47^. cGAS-STING pathway activation has been observed in a cellular model of progeria ^48^ and *Lmna*-knockout developing hearts ^27^. cGAS-STING activation has also been reported in mouse models of other heart diseases ^49–52^. However, a direct link between NE ruptures and the cGAS-STING activation in *LMNA*-related DCM remains undefined. In this study, we report pervasive NE ruptures prior to DCM development and provide evidence against the hypothesis that cGAS-STING activation contributes to *LMNA*-related DCM. Instead, our study nominates a role for extracellular matrix signaling from cardiomyocytes in this disease.

## RESULTS

### Lamin A/C reduction in cardiomyocytes causes dilated cardiomyopathy in adult mice

We generated a mouse model of *LMNA*-related DCM using cardiomyocyte-specific deletion of *Lmna* (*Lmna^F/F^;Myh6-MerCreMer*; *Lmna^CKO^* hereafter) and littermate control wild-type mice (*Lmna^+/+^;Myh6-MerCreMer*) (**Fig. S1A, B**). *Lmna* deletion in cardiomyocytes was induced by tamoxifen administration to adult mice at 6-8 weeks of age (**Fig. 1A**). Lamin A and Lamin C proteins were 47% and 43% reduced, respectively, in cardiomyocytes of *Lmna^CKO^*mice at 2 weeks post tamoxifen, consistent with the 4-week half-life of Lamin A/C in mouse hearts ^53^ (**Fig. 1B; Fig. S1C**). The reduced, but persistent, Lamin A/C in individual cardiomyocytes was confirmed at 1, 2, and 3.5 weeks post tamoxifen (**Fig. 1C**). Thus, *Lmna^CKO^*mice reproduced Lamin A/C protein insufficiency, not elimination, observed in cardiomyocytes of *LMNA*-related DCM patients ^18,20^.

**Figure 1.**
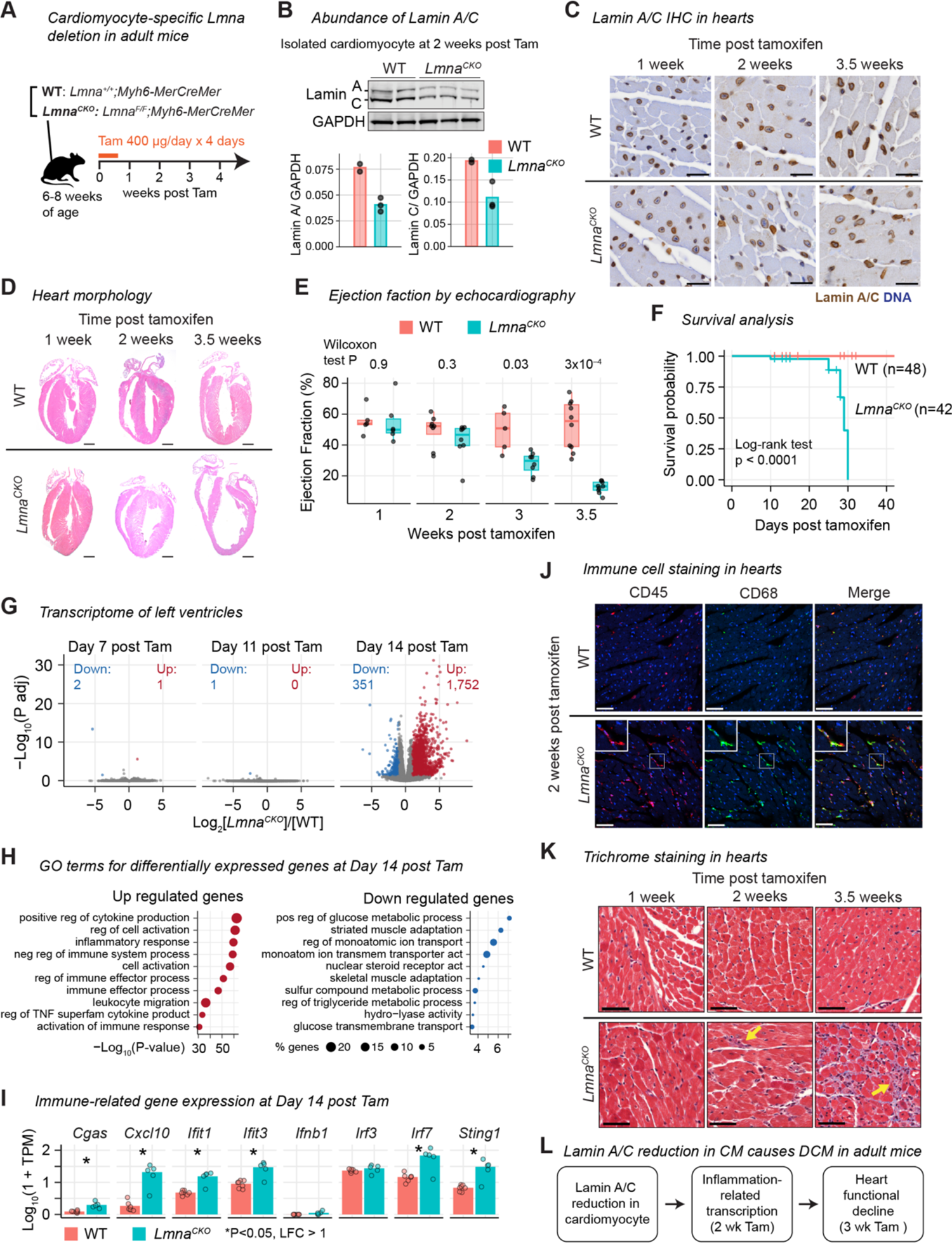
Lamin A/C reduction in cardiomyocytes causes dilated cardiomyopathy in adult mice **A)** Tamoxifen (Tam) induces *Lmna* deletion in cardiomyocytes in adult mice. See **Fig. S1** for related analyses. **B)** Lamin A/C immunoblot in isolated cardiomyocytes (top) with signal quantification (bottom). **C)** Lamin A/C immunohistochemistry (brown) with hematoxylin counterstaining (blue) in mouse heart tissues. Scale bar: 20 μm. **D)** Hematoxylin-eosin staining of hearts. Scale bar: 1 mm. **E)** Left ventricular ejection fraction with interquartile range (box) measured by echocardiography. **F)** Kaplan-Meier survival analysis. **G)** Volcano plot comparing RNA-seq read counts in *Lmna^CKO^* versus WT hearts. Up, upregulated gene count. Down, downregulated gene count. **H)** The top 10 Gene Ontology terms overrepresented among differentially expressed genes. **I)** Normalized RNA-seq gene expressed levels for immune-related genes (bar, mean). **J)** Immunofluorescence of heart tissue sections for CD45 (pan-leukocyte, red) and CD68 (macrophage, green). Scale bar: 20 μm. **K)** Masson’s trichrome staining of heart sections. Arrow, collagen deposition. Scale bar: 20 μm. **L)** Summary

*Lmna^CKO^* mice developed progressive DCM as reported previously (**Fig. 1D, E; Fig. S1D**)^38^. Hearts of *Lmna^CKO^* mice were indistinguishable from wild-type hearts in gross morphology and contractile activity until 2 weeks post tamoxifen. However, at 3 weeks post tamoxifen, left ventricular ejection fraction, an indicator of cardiac systolic function, significantly diminished (**Fig. 1E**). At 3.5 weeks post tamoxifen, *Lmna^CKO^* hearts became severely dilated, with a significant reduction of ventricular wall thickness and loss of systolic activity. *Lmna^CKO^* mice invariably died between 3.5 weeks and 4.5 weeks post tamoxifen (**Fig. 1F**). Thus, the modest reduction of Lamin A/C in cardiomyocytes was sufficient to cause DCM that progressed to heart failure, recapitulating a crucial aspect of human *LMNA*-related DCM.

### Lamin A/C reduction in cardiomyocytes activates inflammation-related transcription

To identify the onset of molecular changes preceding the functional decline of *Lmna^CKO^* hearts, we profiled the transcriptome of the heart at Days 7, 11, and 14 post tamoxifen by RNA-seq (**Table S1**). At Days 7 and 11, the transcriptome of *Lmna^CKO^* hearts was almost identical to that of wild-type hearts (**Fig. 1G**). However, at Day 14, *Lmna^CKO^*hearts exhibited strong transcriptional upregulation (1,751 genes) and modest downregulation (351 genes). The upregulated genes were highly overrepresented for gene ontology (GO) terms related to inflammatory responses (**Fig. 1H**). These upregulated genes included innate immune-related genes such as *Cxcl10*, *Ifit1*, and *Irf7* (**Fig. 1I**), which can be activated by the cGAS-STING pathway. The downregulated genes were overrepresented for gene pathways for cardiomyocyte metabolism and function (**Fig. 1H**).

We examined whether the upregulation of inflammation-related genes reflected an inflammatory response in *Lmna^CKO^* hearts. We observed that the CD45+CD68+ macrophage population increased in *Lmna^CKO^*hearts at 2 weeks post tamoxifen (**Fig. 1J; Fig. S1E**). We also observed increased interstitial collagen deposition at 2 weeks post tamoxifen, which developed into extensive fibrosis by 3.5 weeks post tamoxifen (**Fig. 1K**). Thus, Lamin A/C reduction in cardiomyocytes resulted in extensive upregulation of inflammation-related genes, macrophage expansion, and initial fibrotic response at 2 weeks post tamoxifen, preceding the morphological and functional changes reflective of clinical DCM (**Fig. 1L**).

### *Lmna^CKO^* cardiomyocytes develop pervasive localized nuclear envelope ruptures

We examined whether Lamin A/C reduction caused nuclear envelope (NE) ruptures in cardiomyocytes. Strikingly, we observed protrusion of DNA from one or multiple nuclei inside cardiomyocytes in *Lmna^CKO^*heart sections at 2 weeks post tamoxifen, while this event was absent in wild-type hearts (**Fig. 2A**; of note, 90% of cardiomyocytes are binucleated in adult mice ^54^). Nuclei with protruded DNA were positive for Lamin A/C staining, indicating that partial reduction of Lamin A/C was sufficient to cause DNA protrusion (**Fig. 2A**). Electron microscopy revealed strong condensation of protruded DNA that appeared devoid of the surrounding nuclear membrane, suggesting NE ruptures (**Fig. 2B; Fig. S2A**). We isolated cardiomyocytes from hearts at 2 weeks post tamoxifen, immediately fixed them, and examined the nuclear envelope. Lamin A/C signals were specifically lost at the tips of nuclei from which DNA protruded in *Lmna^CKO^* cardiomyocytes (**Fig. 2C**). The Lamin A/C-lost locations also lost staining for PCM1, a perinuclear matrix protein localized at the cytoplasmic side of the outer nuclear membrane ^55^ (**Fig. 2D**). The loss of Lamin A/C and PCM1 occurred specifically at the tips of the elongated nuclei positioned along the longitudinal axis of the cardiomyocytes (**Fig. 2A–D; Fig. S2A**). The specific loss of nuclear envelope proteins Lamin A/C and PCM1 at nuclear tips suggested localized NE ruptures in *Lmna^CKO^* cardiomyocytes.

**Figure 2.**
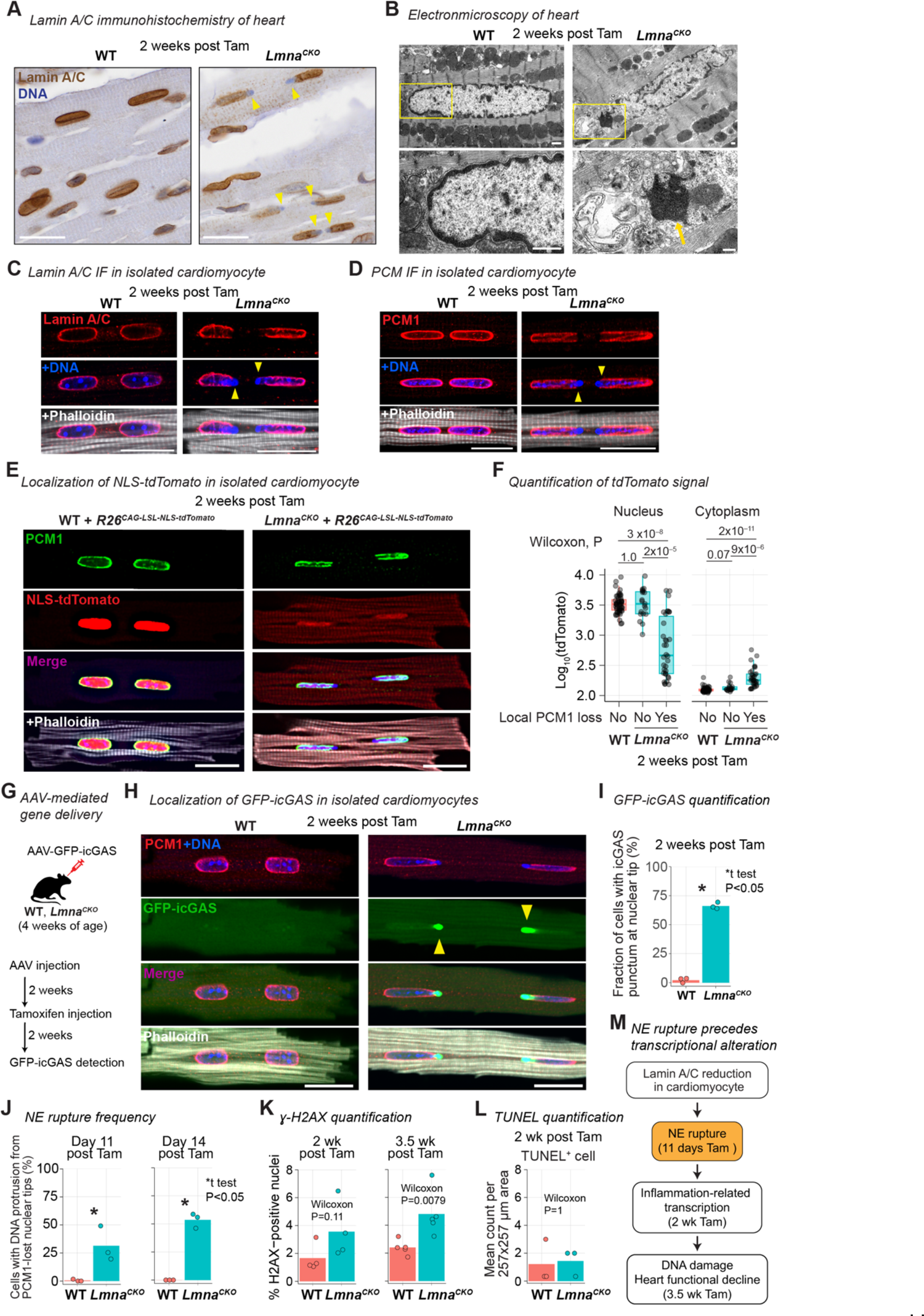
Lamin A/C reduction causes localized nuclear envelope rupture in cardiomyocytes **A)** Lamin A/C immunohistochemistry in heart tissue. Arrow, DNA protruded from nuclei. Scale bar: 20 μm. See **Fig. S2** for related analyses. **B)** Top: Transmission electron micrograph of heart sections focusing on cardiomyocytes. Bottom: Close-up image of the area indicated by rectangle in upper panel. Arrow, protruded chromatin. Scale bar: 1 μm. **C)** Immunofluorescence for Lamin A/C in isolated cardiomyocytes. Phalloidin stains F-actin. Arrowhead, local loss of Lamin A/C with protruded DNA. Scale bar: 20 μm. **D)** Same as **(C)**, but for PCM1. **E)** NLS-tdTomato signals in isolated cardiomyocytes co-stained for PCM-1. Scale bar: 20 μm. **F)** NLS-tdTomato signal intensity in nucleus and cytoplasm of cardiomyocytes (box, interquartile range). *Lmna^CKO^* cardiomyocytes are stratified by the presence or absence of local PCM-1 loss at the nuclear envelope. WT n=51, *Lmna^CKO^* n=53 (PCM1-loss 34, intact 19) cardiomyocytes. **G)** MyoAAV-mediated GFP-icGAS expression in cardiomyocytes *in vivo*. **H)** GFP-icGAS signals in isolated cardiomyocytes co-stained for PCM1. Scale bar: 20 μm. **I)** Percentage of cardiomyocytes with GFP-icGAS punctum at nuclear tip (bar, mean). **J)** Percentage of cardiomyocytes with DNA protrusion from PCM1-lost nuclear tip (bar, mean) at 11 days (left) and 14 days (right) post tamoxifen. **K)** Percentage of gamma-H2AX-positive nuclei in heart section (bar, mean). **L)** Number of TUNEL-positive cells in heart section (bar, mean). **M)** Summary

To determine whether NE ruptures occurred in *Lmna^CKO^*cardiomyocytes, we examined the retention of nuclear-localized tdTomato. We expressed nuclear localization signal (NLS)-fused tdTomato in cardiomyocytes from a transgene *in vivo* and investigated tdTomato localization immediately after cardiomyocyte isolation at 2 weeks post tamoxifen. NLS-tdTomato was localized exclusively to the nucleus in wild-type cardiomyocytes as well as to the intact nuclei in *Lmna^CKO^*cardiomyocytes (**Fig. 2E; Fig. S2B**). However, NLS-tdTomato signals significantly diminished in nuclei with local PCM1 loss in *Lmna^CKO^* cardiomyocytes (**Fig. 2E, F**). Reciprocally, NLS-tdTomato signals increased in the cytoplasm of *Lmna^CKO^* cardiomyocytes with PCM1-lost nuclei. NLS-tdTomato leakage from damaged nuclei indicated that the nuclear envelope had ruptured in *Lmna^CKO^* cardiomyocytes.

To quantify NE rupture events, we expressed GFP-tagged catalytically-inactive cGAS (icGAS) in cardiomyocytes *in vivo* using muscle-tropic adeno-associated virus (MyoAAV) ^56^ (**Fig. 2G**). icGAS is widely used as a “NE rupture marker” owing to the cGAS’s affinity to cytoplasmic DNA ^57,58^. We observed strong icGAS punctum at every PCM1-lost site on the nuclear envelope in *Lmna^CKO^* cardiomyocytes at 2 weeks post tamoxifen (**Fig. 2H; Fig. S2C**). We found that 66% of *Lmna^CKO^* cardiomyocytes have at least one nucleus with icGAS puncta on nuclear envelopes, while icGAS puncta were virtually absent in wild-type cardiomyocytes (**Fig. 2I**). Thus, NE ruptures were pervasive in *Lmna^CKO^* cardiomyocytes at 2 weeks post tamoxifen.

To determine whether NE ruptures preceded the transcriptional change, we quantified NE ruptures at Day 11 post tamoxifen, when no transcriptional change was detected (**Fig. 1G**). We used a local loss of PCM1 with concomitant cytoplasmic DNA protrusion as a proxy for NE ruptures, as this feature strongly correlated with icGAS localization. We observed that 31% of *Lmna^CKO^* cardiomyocytes had at least one ruptured nucleus at Day 11 (**Fig. 2J; Fig. S2D, E**). At Day 14, the NE rupture frequency was 54% based on this method, which was comparable to 66% based on the sensitive icGAS-based quantification (**Fig. 2J**). Thus, about one third of *Lmna^CKO^* cardiomyocytes developed NE ruptures as early as Day 11 post tamoxifen, preceding the earliest transcriptional changes, with more than 50% of nuclei presenting ruptures by Day 14.

We investigated whether the pervasive NE ruptures in *Lmna^CKO^*cardiomyocytes accompanied DNA damage accumulation (**Fig. 2K; Fig. S2F**). We did not find a statistically significant increase of gamma-H2AX-positive nuclei indicative of DNA double-strand break in *Lmna^CKO^* hearts at 2 weeks post tamoxifen. In addition, there was no indication of increased cell death assessed by TUNEL at this time point (**Fig. 2L; Fig. S2G**). However, at 3.5 weeks post tamoxifen, gamma-H2AX-positive nuclei significantly increased. Thus, pervasive NE ruptures did not cause immediate DNA damage accumulation or cell death at the time of strong inflammation-related transcriptional upregulation. Instead, DNA damage accumulated when the heart structure and function began to deteriorate (**Fig. 2M**).

### *Lmna^CKO^* cardiomyocytes do not activate cGAS-STING-related transcription

Given the pervasive NE ruptures, we tested whether the cGAS-STING cytosolic DNA sensing pathway was activated within *Lmna^CKO^* cardiomyocytes. In the cGAS-STING pathway, cGAS detection of cytoplasmic DNA activates STING-mediated interferon transcription within cells bearing the cytoplasmic DNA ^46,47^. This pathway has been implicated in Lamin A/C-based cardiomyopathy in a previous study ^27^. cGAS and STING proteins were expressed in the heart, but interestingly, their expression within cardiomyocytes was very low in both wild-type and *Lmna^CKO^* backgrounds (**Fig. 3A; Fig. S3A, B**). To test whether cGAS-STING-downstream genes were upregulated within cardiomyocytes of *Lmna^CKO^*hearts, we distinguished cardiomyocyte and non-cardiomyocyte transcriptomes in the heart using SLAM-IT-seq^59^. SLAM-IT-seq labels transcripts in *Cre*-positive cardiomyocytes with 4-thiouracil (4sU) and distinguishes them from unlabeled transcripts from non-cardiomyocyte cells (**Fig. S3C**). As expected, cardiomyocyte-specific genes were specifically 4sU-labeled in wild-type hearts (**Fig. S3D**). We then assessed the labeling state of the 1,020 upregulated genes in *Lmna^CKO^* hearts and found that 135 genes were 4sU-labeled (i.e. cardiomyocyte-derived) and 885 genes were not 4sU-labeled (i.e. non-cardiomyocyte-derived) in *Lmna^CKO^* hearts at 2 weeks post tamoxifen (**Fig. 3B**). The cardiomyocyte-originated upregulated genes were not overrepresented for inflammation-related GO terms (**Fig. 3B**) and did not include well-established cGAS-STING downstream genes (**Fig. S3E**; **Table S2**). Instead, these genes were overrepresented for the ECM-related GO terms such as “Proteoglycans in cancer”. On the other hand, non-cardiomyocyte-originated upregulated transcripts were strongly enriched for inflammation-related GO terms (**Fig. 3B**). These data suggested that the cGAS-STING pathway was not activated within *Lmna^CKO^*cardiomyocytes and that the inflammation-related transcriptional activation originated from non-cardiomyocytes in *Lmna^CKO^* hearts.

**Figure 3.**
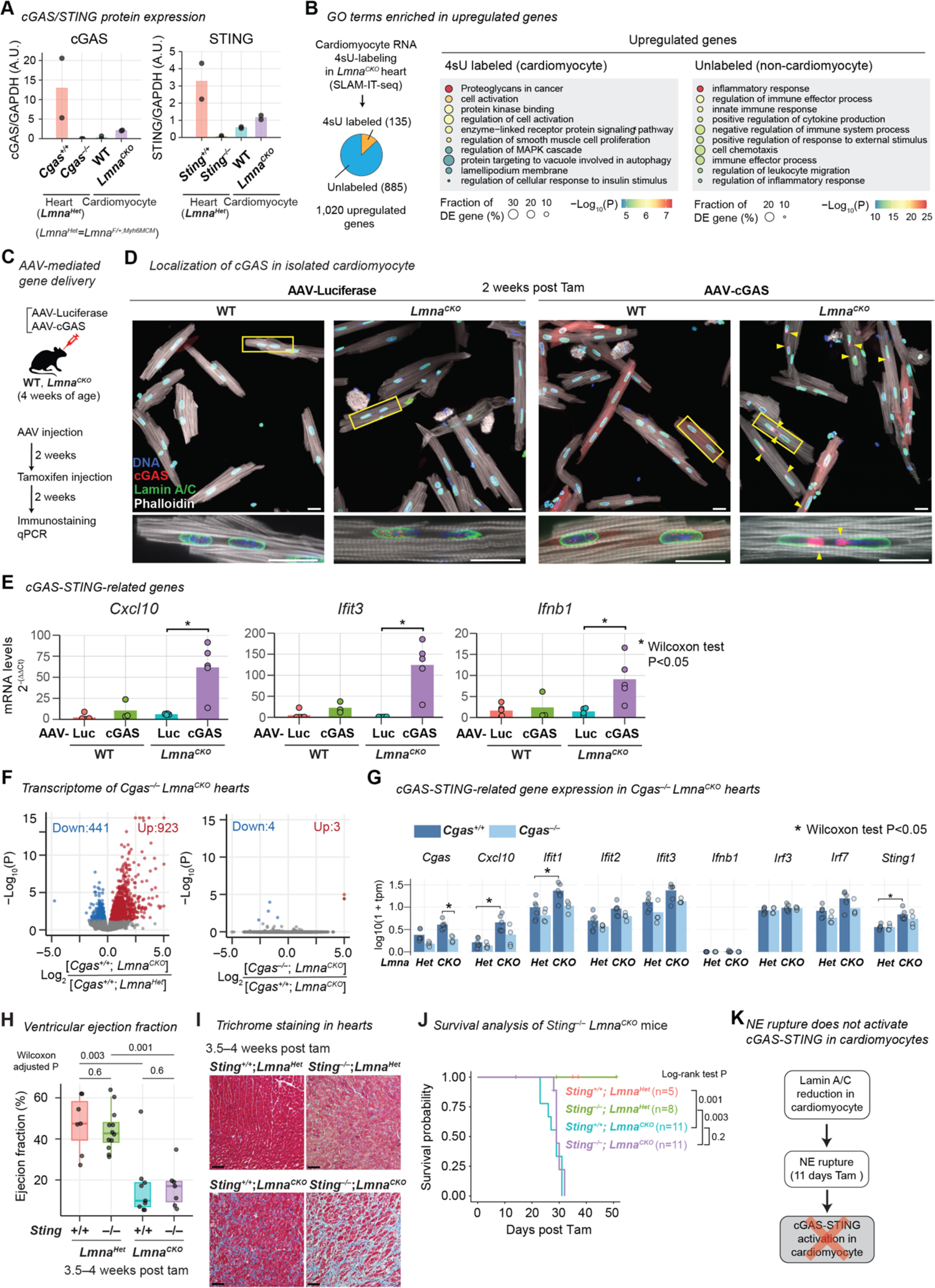
*Cgas* and *Sting* are not required for inflammation-related gene expression and dilated cardiomyopathy in *Lmna^CKO^* mice **A)** Normalized cGAS and STING abundance quantified by immunoblot shown in **Fig. S3A, B** (bar, mean). **B)** Left: Upregulated genes in *Lmna^CKO^* hearts at 2 weeks post tamoxifen are categorized into cardiomyocyte-derived or non-cardiomyocyte-derived based on 4sU labeling in SLAM-IT-seq. Right: GO terms enriched among each class of upregulated genes. **C)** MyoAAV-mediated GAS overexpression in cardiomyocytes *in vivo*. **D)** Top: Anti-cGAS and Lamin A/C immunofluorescence in isolated cardiomyocytes. Bottom: Close-up image of the area indicated by rectangle in upper panel. Arrowhead, GFP-icGAS localization at nuclear tip. Scale bar: 20 μm. **E)** Transcript abundance in isolated cardiomyocytes quantified by quantitative PCR and normalized by *Actb* transcripts (bar, mean). **F)** Volcano plot comparing RNA-seq read counts in *Cgas^-/-^*; *Lmna^Het^* (n=4) versus *Cgas^+/+^*; *Lmna^Het^* (n=6) (left) and in *Cgas^-/-^*; *Lmna^CKO^* (n=4) versus *Cgas^+/+^*; *Lmna^CKO^*(n=6) (right). **G)** Normalized RNA-seq gene expressed levels for cGAS-STING-related genes (bar, mean). **H)** Left ventricular ejection fraction measured by echocardiography (box, interquartile range). **I)** Masson’s trichrome staining of heart sections. Scale bar: 20 μm. **J)** Kaplan-Meier survival analysis. **K)** Summary

We hypothesized that the near absence of cGAS expression prevented cGAS-STING activation within *Lmna^CKO^* cardiomyocytes. We tested this hypothesis by overexpressing wild-type cGAS in cardiomyocytes *in vivo* with MyoAAV (**Fig. 3C**). We first confirmed that endogenous cGAS was not detectable at NE rupture sites (**Fig. 3D**) and that the well-established cGAS-STING downstream genes, *Cxcl10*, *Ifit3*, and *Ifnb1* ^60–63^, remained transcriptionally silent in cardiomyocytes of *Lmna^CKO^* mice transduced with control luciferase MyoAAV (**Fig. 3E**). However, cGAS overexpression in *Lmna^CKO^* cardiomyocytes resulted in strong cGAS accumulation at NE rupture sites (**Fig. 3D**). Consistently, *Cxcl10*, *Ifit3*, and *Ifnb1* transcripts were 10-, 128-, and 6-fold upregulated, respectively, upon cGAS overexpression (**Fig. 3E**). In wild-type mice, cGAS overexpression did not upregulate cGAS-downstream genes, confirming the NE rupture-dependency of cGAS-downstream activation upon cGAS overexpression. Taken together, these data suggested that *Lmna^CKO^* cardiomyocytes did not activate the cGAS-STING pathway despite pervasive NE ruptures and that the lack of cGAS-STING activation was likely due to the low cGAS expression in adult cardiomyocytes.

### *Cgas* and *Sting* are not required for inflammation-related gene expression and DCM in *Lmna^CKO^* mice

We examined the genetic requirement for *Cgas* and *Sting*, the essential mediators of the cGAS-STING pathway, to the inflammation-related transcriptional activation and DCM in *Lmna^CKO^* hearts. We introduced the germ-line *Cgas^-/-^* allele ^64^ into *Lmna^CKO^* mice or control *Lmna^Het^* mice (*Lmna^F/+^;Myh6MerCreMer*) and validated the absence of cGAS protein in the heart of *Cgas^-/-^*carriers (**Fig. S3A**). We then performed RNA-seq in whole hearts at 2 weeks post tamoxifen. We first confirmed the expected strong upregulation of inflammation-related genes in *Cgas^+/+^*;*Lmna^CKO^* hearts compared with *Cgas^+/+^*;*Lmna^Het^* hearts (**Fig. 3F; Fig. S3F; Table S1**). We then compared *Cgas^-/-^*;*Lmna^CKO^*hearts with *Cgas^+/+^*;*Lmna^CKO^* hearts. There was almost no difference in gene expression between them (**Fig. 3F**). Representative cGAS-STING-downstream genes did not show statistically significant differences in gene expression between *Cgas^-/-^*;*Lmna^CKO^* and *Cgas^+/+^*;*Lmna^CKO^* hearts (**Fig. 3G; Fig. S3G**). These data suggested that the inflammation-related transcription in *Lmna^CKO^*hearts was independent of cGAS.

We next combined germ-line *Sting^-/-^* mice ^65^ with *Lmna^CKO^* mice (**Fig. S3B**) and investigated the heart phenotype. By 3.5 to 4 weeks post tamoxifen, *Sting^-/-^*;*Lmna^CKO^* mice showed as severely reduced left ventricular ejection fraction (**Fig. 3H**) and extensive fibrosis (**Fig. 3I**) as the littermate control *Sting^+/+^*;*Lmna^CKO^* mice did, whereas *Sting^+/+^*;*Lmna^Het^* and *Sting^-/-^*;*Lmna^Het^* mice maintained normal hearts. Likewise, *Cgas^-/-^*;*Lmna^CKO^* hearts developed severe fibrosis (**Fig. S3H**). Moreover, both *Sting^-/-^*;*Lmna^CKO^* mice and *Cgas^-/-^*;*Lmna^CKO^* mice died as early as *Sting^+/+^*;*Lmna^CKO^* or *Cgas^+/+^*;*Lmna^CKO^* mice did, whereas no lethality was observed in *Sting^-/-^*;*Lmna^Het^* and *Cgas^-/-^*;*Lmna^Het^*mice (**Fig. 3J; Fig. S3I**). Taken together, our data suggested that the cGAS-STING pathway did not mediate inflammatory transcription and the pathophysiologic features of DCM in *Lmna^CKO^* hearts (**Fig. 3K**).

### Cardiomyocytes, fibroblasts, and immune cells are transcriptionally altered in *Lmna^CKO^* hearts

We explored alternative mechanisms by which Lamin A/C reduction in cardiomyocytes caused DCM. To identify intercellular signaling initiated by *Lmna^CKO^* cardiomyocytes, we defined the cell type-resolved transcriptional changes using single-nucleus (sn) RNA-seq (**Table S3**). We obtained transcriptomes for 14,111 nuclei from either wild-type or *Lmna^CKO^* hearts at 2 weeks post tamoxifen (n=3; **Fig. 4A**). These nuclei were grouped into cardiomyocytes (CM1 and CM2), fibroblasts (Fib1 and Fib2), macrophages (Mac1 and Mac2), T cells, endothelial cells (coronary, lymphatic, and other), pericytes, or neurons, based on their transcriptomes (**Fig. S4A**). Among these, CM2, Fib2, Mac2, and T-cells were highly overrepresented by *Lmna^CKO^* heart-derived cells (**Fig. 4B; Fig. S4B**). CM2 was characterized by high *Rtn4* expression, a feature also reported in *Lmna*-knockout myotubes ^66^ and Lamin A/C-independent heart disease ^67–70^ (**Fig. S4A**). Fib2 was characterized by myofibroblast marker *Postn* ^71^, Mac2 by circulating monocytic receptor *Ccr2* ^72^, and T cells by T cell adaptor *Skap1* ^73^. We found extensive differential gene expression within cardiomyocytes (CM1 and CM2 combined), fibroblasts (Fib1 and Fib2 combined), and immune cells (Mac1, Mac2, T cell combined) while other cell types exhibited almost no expression differences (**Fig. 4C, D; Fig. S4C; Table S4**). The upregulated genes in cardiomyocytes were not overrepresented for cytosolic DNA sensing pathway, did not include cell death-related genes, but included some DNA damage repair-related genes, as expected (**Fig. S4D–F**).

**Figure 4.**
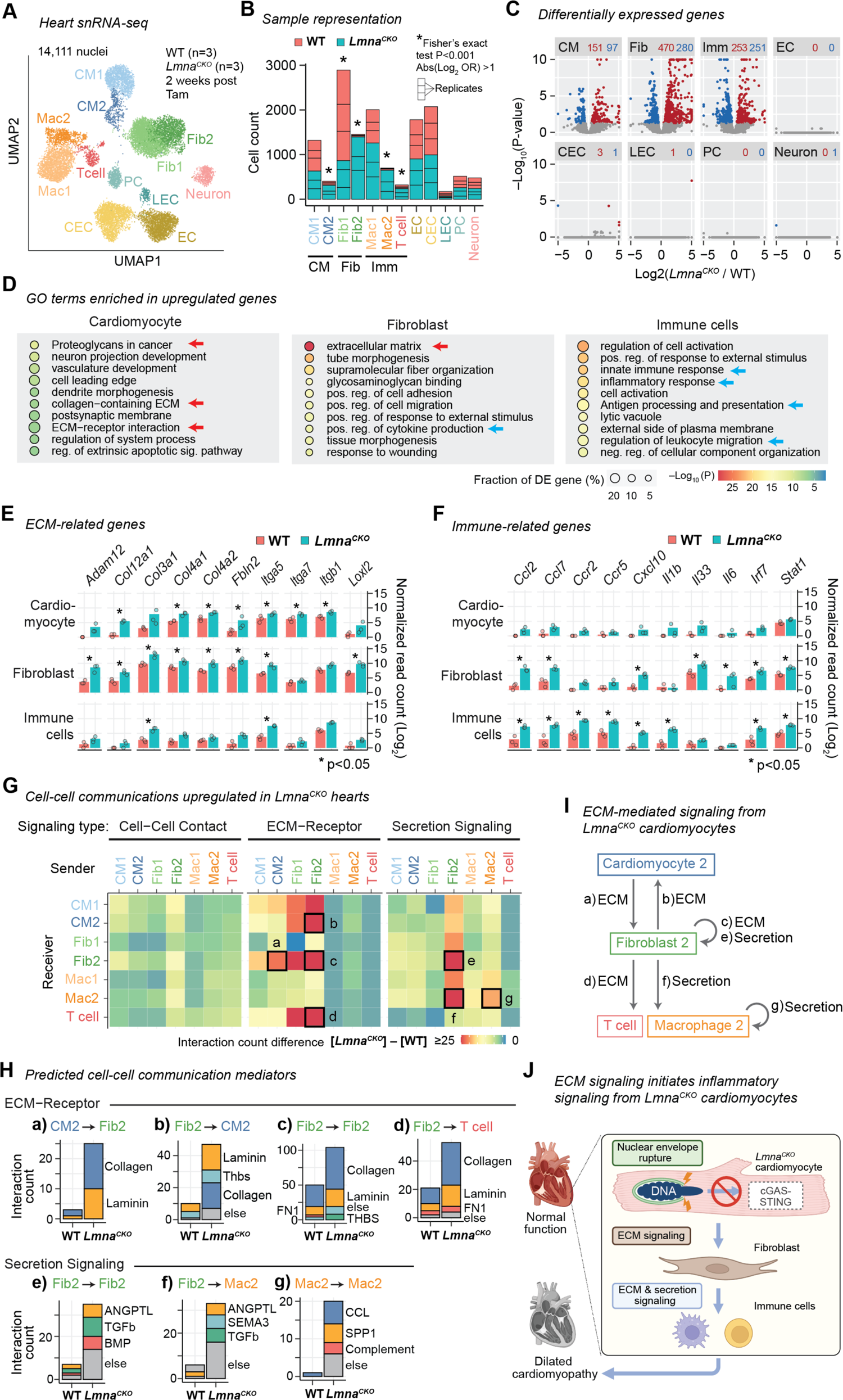
ECM-mediated signaling from *Lmna^CKO^* cardiomyocytes is predicted to activate fibroblasts **A)** snRNA-seq UMAP plot for 14,111 nuclei from either WT or *Lmna^CKO^* hearts, colored by cell-type annotation. CM, cardiomyocyte. Fib, fibroblast. Mac, macrophage. EC, endothelial cell. CEC, coronary EC. LEC, lymphatic EC. PC, pericyte. See **Fig. S4** for additional analyses. **B)** Count of WT and *Lmna^CKO^* heart-originating nuclei within each cell type cluster. Cell types combined in later analyses are indicated below graph. **C)** Pseudo-bulk volcano plot comparing sbRNA-seq read counts in cells of *Lmna^CKO^*hearts versus cells of WT hearts within each cell type. Red, upregulated genes. Blue, downregulated genes. **D)** GO terms enriched among upregulated genes within indicated cell type. Red arrows, ECM-related GO term. Blue arrow, inflammation-related GO term. **E)** Pseudo-bulk transcript abundance of ECM-related genes. Point, mean of normalized read count per replicate. Bar, mean across replicates. **F)** Same as (E), but for immune-related genes. **G)** Predicted signaling strength between sender cells (x-axis) and receiver cells (y-axis). Color indicates the extent of signaling gains in *Lmna^CKO^* hearts relative to WT hearts. **H)** Predicted signaling mediators for the signaling indicated by box in (**G**) with alphabetical labels. **I)** Predicted intercellular signaling (indicated in **G** by alphabet labels) in *Lmna^CKO^* hearts. **J)** Model

Interestingly, upregulated genes in *Lmna^CKO^* cardiomyocytes and those in fibroblasts were both most strongly overrepresented for ECM-related pathways (**Fig. 4D**). The upregulated ECM-related genes included collagen and integrin genes (in cardiomyocytes and fibroblasts) and ECM proteinase and crosslinker genes (specifically in fibroblasts) (**Fig. 4E**). On the other hand, fibroblasts and immune cells, but not cardiomyocytes, commonly upregulated immune-related genes including cytokine, cytokine receptor, and cytokine transcription factor genes (**Fig. 4D, F**). These results suggested potential signal crosstalk between cardiomyocytes and fibroblasts and between fibroblasts and immune cells in *Lmna^CKO^* hearts.

### ECM-mediated signaling from *Lmna^CKO^* cardiomyocytes is predicted to activate fibroblasts

We investigated intercellular signal crosstalk in *Lmna^CKO^*hearts by applying the CellChat program ^74^ to the snRNA-seq data. This analysis predicted ECM-receptor signaling from CM2 to Fib2 as the only strongly upregulated signaling originating from CM2, the population predominantly consisted of *Lmna^CKO^*cardiomyocyte nuclei (**Fig. 4G, Box a**). Collagens and laminins were predicted to mediate this CM2-to-Fib2 signaling (**Fig. 4H**). CM2 were not predicted to signal directly to immune cell populations. Instead, Fib2 were predicted to send ECM-mediated signals to CM2 (**Box b**), Fib2 (**Box c**), and T cells (**Box d**), via collagens, laminins, thrombospondins, and fibronectins (**Fig. 4G, H**). Fib2 were also predicted to send secretion-mediated signals to Fib2 (**Box e**), and Mac2 (**Box f**), via ANGPTL (angiopoietin-like), TGF beta, BMP, and SEMA3 (semaphorin3). Fib2 might therefore act as a central signaling hub, receiving signals from CM2 and sending signals to immune cells (**Fig. 4I**). Finally, Mac2 were predicted to send secretion-mediated signals to Mac2 themselves (**Box g**) via CCL (chemokine ligands), SPP1 (osteopontins), and complement factors. This signaling might mediate the macrophage infiltration observed in *Lmna^CKO^* hearts (**Fig. 1J**). Taken together, our results suggested that *Lmna^CKO^* cardiomyocytes activated fibroblasts via ECM-mediated signaling, not cytokine-mediated signaling such as cGAS-STING, and that fibroblasts activated and recruited immune cells through both ECM signaling and secretion signaling to orchestrate inflammatory signaling in the heart (**Fig. 4J**).

## DISCUSSION

We report that frequent localized NE ruptures precede transcriptional changes, DNA damage accumulation, and heart functional decline in *Lmna^CKO^* hearts. Given our observation that a 50% reduction of Lamin A/C proteins is sufficient to induce pervasive NE ruptures in cardiomyocytes, NE ruptures might be more prevalent than previously anticipated in *LMNA*-related DCM. NE ruptures might have been overlooked due to the specific location of localized NE ruptures at the tips of elongated nuclei within cardiomyocytes. The mechanisms underlying NE ruptures and the causal role of NE ruptures in DCM remain to be investigated.

Our data indicate that the cGAS-STING pathway does not contribute to *LMNA*-related DCM in the adult mouse model investigated herein. This suggests that pharmacologically reducing cGAS-STING activity for treating *LMNA*-related DCM, an idea previously discussed ^27,75^, may be ineffective. Our conclusion diverges from a previous report that *Cgas* deletion can delay early postnatal cardiomyopathy caused by *Lmna* deletion in embryonic hearts ^27^. This difference may be due to that the near absence of cGAS expression prevents cGAS-STING activation in adult cardiomyocytes, whereas the basal cGAS-STING activity may be higher in early postnatal cardiomyocytes ^76,77^. The lack of robust cGAS expression may reduce pathogen-derived DNA sensing in adult cardiomyocytes, similar to hepatocytes that also lack functional cGAS-STING ^78^. Many reports suggest an association between DNA damage and cGAS-STING activation ^27,79–82^. Our data that *Lmna^CKO^* cardiomyocytes accumulate DNA damage without activating cGAS-STING suggest that DNA damage and cGAS-STING can be uncoupled.

Our study nominates ECM signaling as a mechanism by which Lamin A/C-reduced cardiomyocytes initiate an inflammatory response by activating fibroblasts. A recent study reported a similar observation in which NE rupture induces ECM remodeling through unknown mechanisms that require DNA damage ^83^. Further study on the potential roles of ECM signaling in *LMNA*-related DCM is required.

### Limitations of the study

We reduced Lamin A/C exclusively within the cardiomyocytes of adult mice. Whether Lamin A/C reduction in other cell types or at other stages causes NE ruptures and contributes to DCM through cGAS-STING activation remains an open question.

## ACKNOWLEDGEMENT

We thank core facilities for animal care, microscopy, genomics, and pathology at Cincinnati Children’s and the University of Chicago for their assistance. This work is supported by NIH grant R21/R33 AG054770 (K.I.), Cincinnati Children’s Research Innovation and Pilot grant (K.I.), NIH grants R01 HL163523, R01 HL124836, and R01 HL126509 (I.P.M).

## AUTHOR CONTRIBUTION

Conceptualization, A.E. and K.I.; Methodology, A.E., I.P.M., and K.I.; Formal Analysis, A.E. and K.I.; Investigation, A.E., H.B., B.T., A.S., S.I., B.L., O.A., P.P., K.I.; Resources, I.P.M.; Data Curation, A.E. and K.I; Writing – Original Draft, A.E., and K.I.; Writing – Review & Editing, A.E., I.P.M., and K.I.; Visualization, A.E. and K.I.; Supervision, K.I.; Project Administration, K.I.; Funding Acquisition, K.I. and I.P.M.

## SUPPLEMENTARY FIGURE LEGENDS

**Figure S1.**
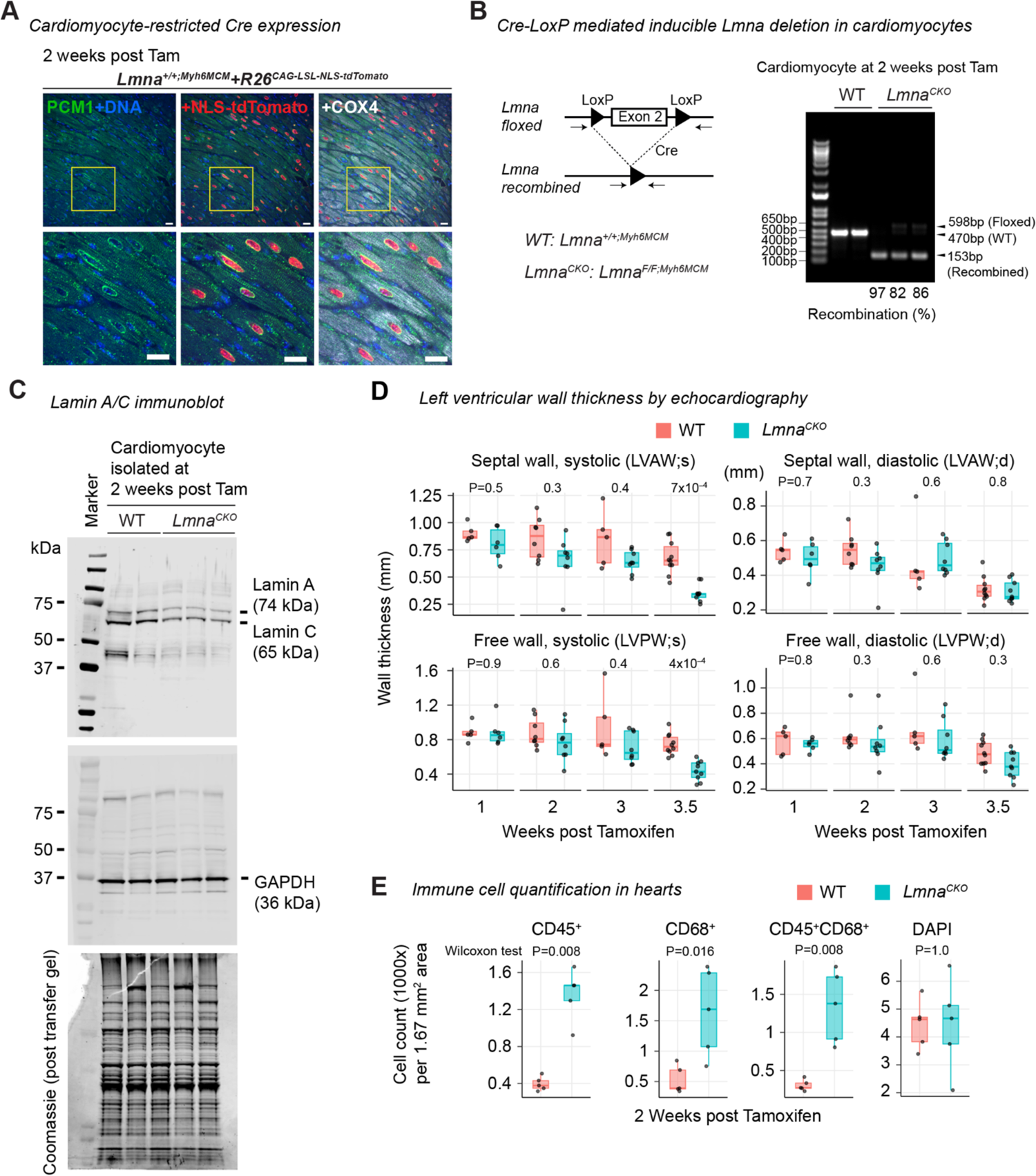
Characterization of adult mice with cardiomyocyte-specific *Lmna* deletion **A)** Immunofluorescence for PCM1, NLS-tdTomato, and COX4 in heart sections. PCM1 and COX4 are used to stain cardiomyocyte nuclei and the cytoplasm, respectively. Scale bar: 20 μm. **B)** Left: Schematic for the *Lmna* LoxP allele and the expected recombined allele upon Cre expression. Arrow, PCR primer location. Right: Amplification of the wild-type, floxed, and recombined *Lmna* alleles by PCR using primers indicated in the left panel. The recombination efficiency (%) was calculated by the intensity of the 153-bp recombined band over the intensity of the 598-bp floxed band. **C)** The whole immunoblot for Fig. 1B, with Coomassie staining of the original gel. **D)** Echocardiography of left ventricular (LV) geometry. P, Wilcoxon test. **E)** Quantification of CD45 and CD68-positive cells in heart sections. Representative image in Fig. 1J.

**Figure S2.**
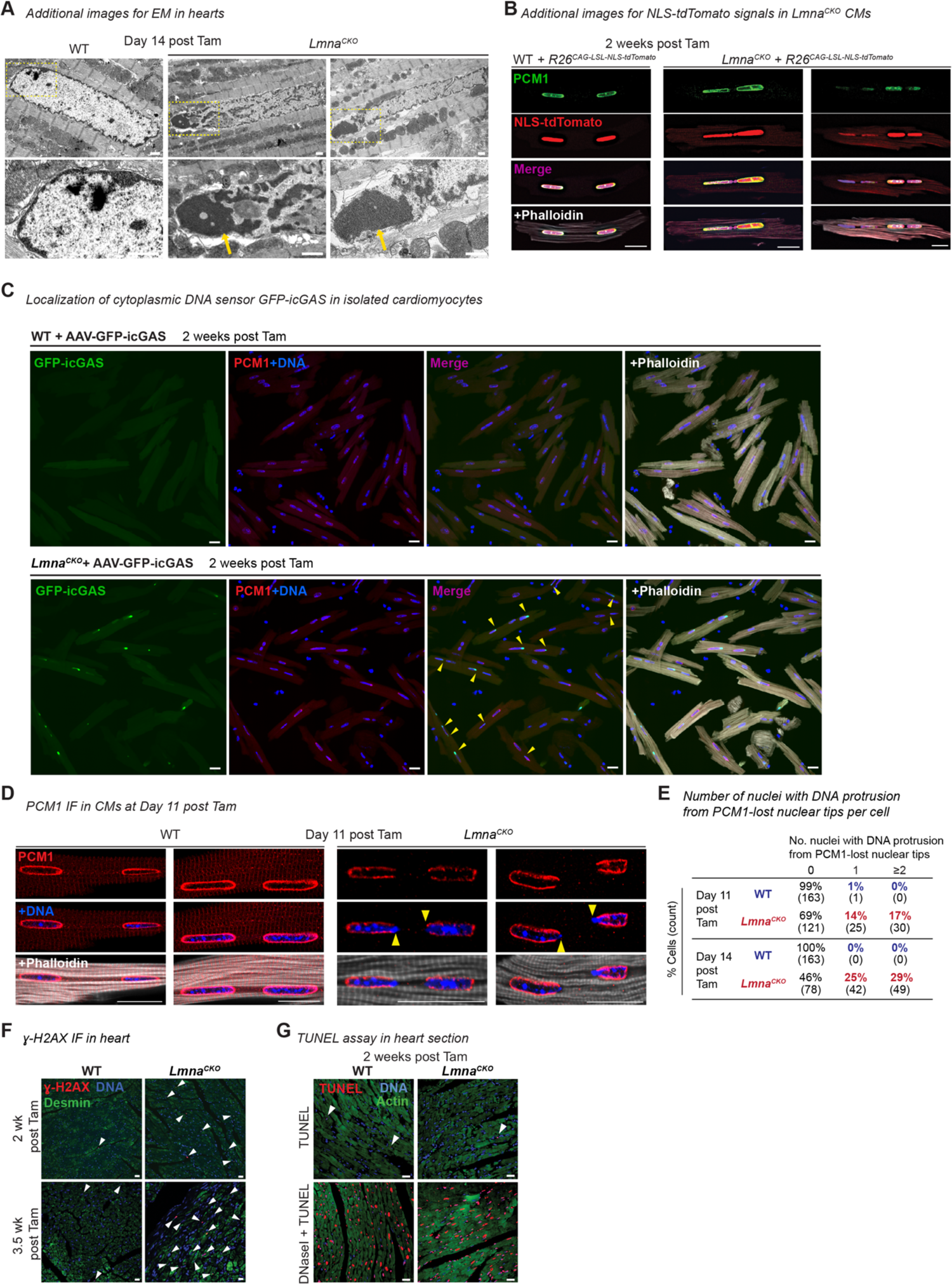
Characterization of nuclear envelope ruptures in *Lmna^CKO^* cardiomyocytes **A)** Top: Additional transmission electron micrographs of heart sections focusing on cardiomyocytes, related to Fig. 2B. Bottom: Enlarged images of the area indicated by yellow box in the upper panels. Arrow, protruded chromatin. Scale bar: 1 μm. **B)** Additional immunofluorescence images for PCM-1 with detection of native NLS-tdTomato. Scale bar: 20 μm. **C)** Immunofluorescence for PCM-1 with detection of exogenous GFP-icGAS in isolated cardiomyocytes, related to Fig. 2H. Phalloidin stains F-actin. Arrowhead, GFP-icGAS localization at nuclear tips. Scale bar: 20 μm. **D)** Immunofluorescence images for PCM1 in isolated cardiomyocytes at Day 11 post tamoxifen. Arrowhead, local loss of PCM1 at nuclear tips. Scale bar: 20 μm. **E)** Fraction of cardiomyocytes with DNA protrusion from PCM1-lost nuclear tips. Cardiomyocytes are stratified by the number of ruptured nuclei per cell. Only multinucleated cardiomyocytes are analyzed. **F)** Immunofluorescence for γH2AX and desmin in heart tissue sections. Desmin stains cardiomyocytes. Arrowhead, γH2AX-stained nuclei. Scale bar: 20 μm. **G)** TUNEL assay for cell death detection in heart tissues. Arrowhead, TUNEL-positive cells. Scale bar: 20 μm.

**Figure S3.**
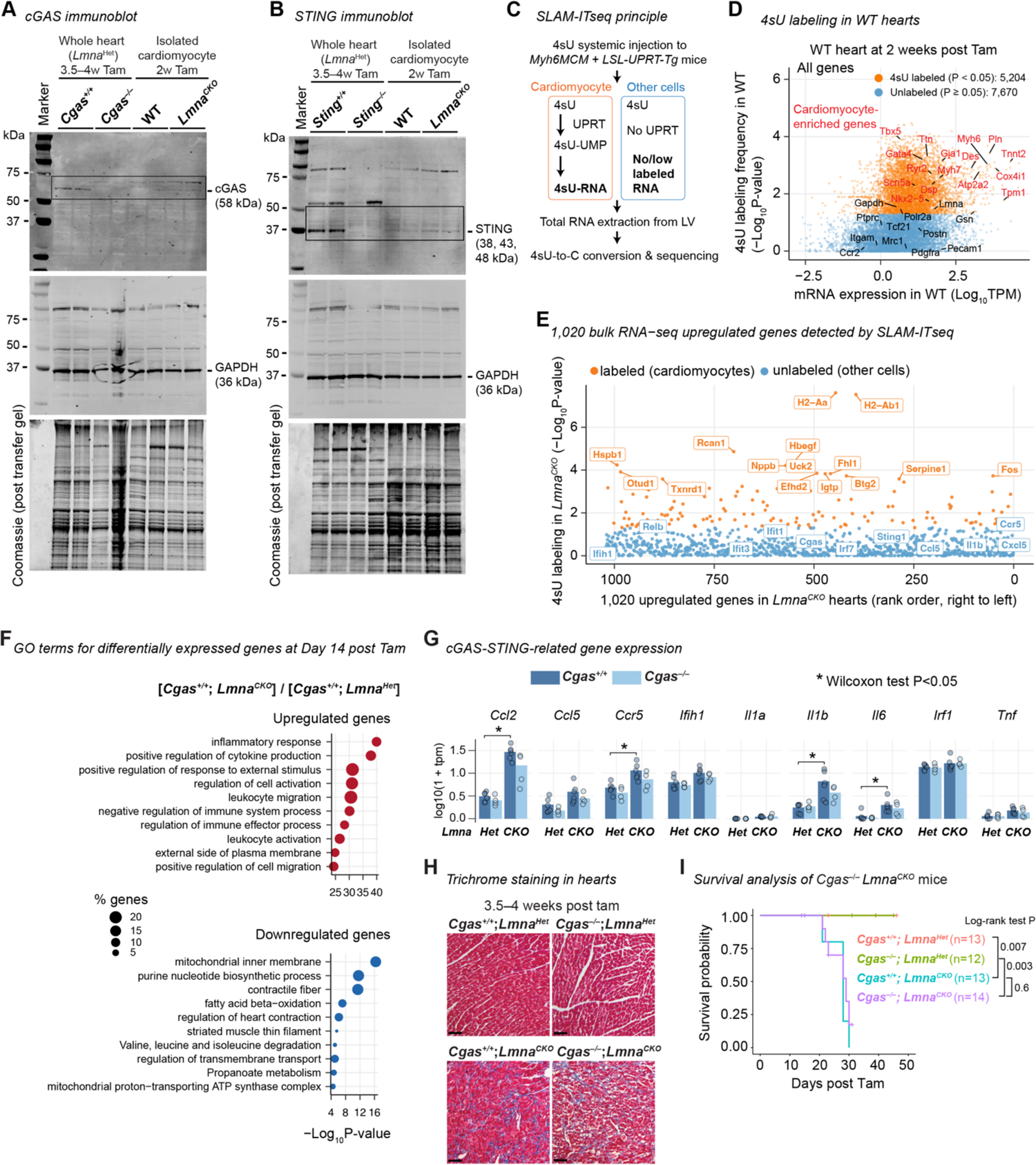
Analysis of the cGAS-STING pathway in *Lmna^CKO^* hearts **A)** cGAS and GAPDH immunoblots, and Coomassie staining of original gel, in hearts and isolated cardiomyocytes, related to Fig. 3A. **B)** Same as (A), but for STING. **C)** Principle of SLAM-IT-seq. Transcripts in cardiomyocytes are labeled by 4-thiouracil (4sU). **D)** SLAM-IT-seq data for all genes in WT hearts at 2 weeks post tamoxifen, with the expression level on the X axis and the 4sU labeling frequency on the Y axis. Select genes known to be preferentially expressed in cardiomyocytes (red) and those not known to be preferentially expressed in cardiomyocytes (black) are indicated. **E)** SLAM-IT-seq data for the 1,020 upregulated genes at 2 weeks post tamoxifen, with the rank order of the fold change of upregulation on the X axis and the 4sU labeling frequency in *Lmna^CKO^* hearts on the Y axis. All cytokine genes among the upregulated genes, which are all among unlabeled genes (blue), are indicated. Highly labeled genes (Log_10_ P-value > 3.5) are also indicated (orange). **F)** The top 10 Gene Ontology terms overrepresented among differentially expressed genes in *Cgas^-/-^*; *Lmna^Het^* versus *Cgas^+/+^*; *Lmna^Het^*. **G)** Normalized RNA-seq gene expressed levels for additional cGAS-STING-related genes (bar, mean). **H)** Masson’s trichrome staining of heart sections. Scale bar: 20 μm. **I)** Kaplan-Meier survival analysis.

**Figure S4.**
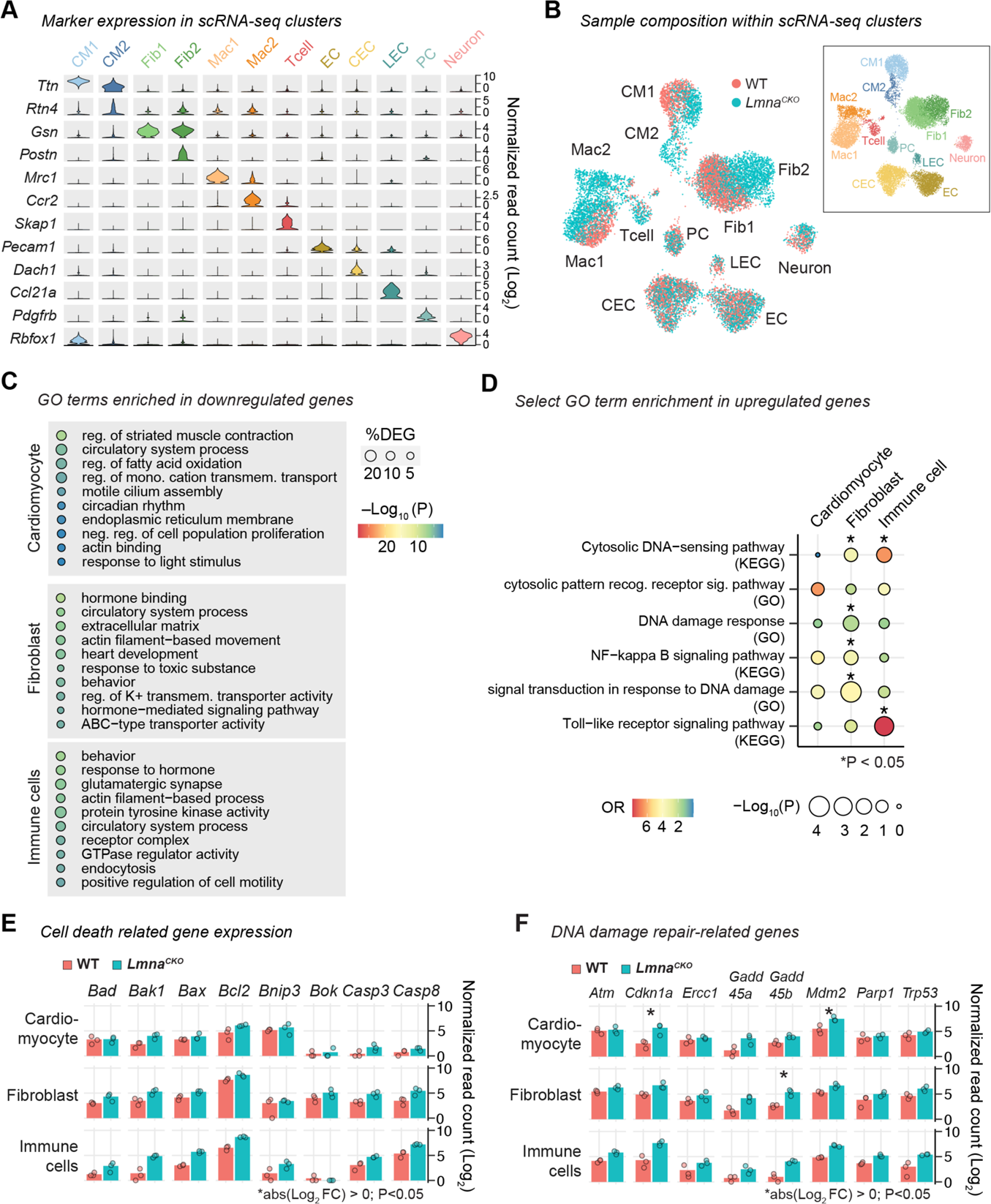
Single-nucleus RNA-seq analysis of *Lmna^CKO^* hearts **A)** Expression (sum of single-nucleus normalized read count across 3 mice within cell type) of maker genes used to classify single nuclei to the cell types indicated along the Y axis, related to Fig. 4A**. B)** snRNA-seq UMAP plot for 14,111 nuclei, colored by sample genotype. Inset, UMAP colored by cell type for reference. **C)** GO terms enriched among downregulated genes within indicated cell type. **D)** Enrichment for select GO terms relevant to cytosolic DNA sensing among upregulated genes. **E)** Expression (mean of single-nucleus normalized read count within cell type) of cell death-related genes. Dot, mean within individual mice. Bar, mean across three mice. **F)** Same as (C), but for DNA damage repair-related genes.

## SUPPLEMENTARY TABLES

**Table S1.** Differentially expressed genes identified by bulk RNA-seq, related to **Figures 1 and 3**. The table includes differentially expressed genes between wild-type and *Lmna^CKO^* hearts at 1 week post tamoxifen, 11 days post tamoxifen, and 2 weeks post tamoxifen, and for *Cgas/Lmna* double-knockout experiment.

**Table S2.** Normalized expression values, C-to-T conversion rates, and beta-binomial test P-values for SLAM-IT-seq data, related to **Figure 3**.

**Table S3.** Indexed oligonucleotides used in snRNA-seq, related to **Figure 4**, including indexed RT primers, indexed ligation primers, and indexed P7 PCR primers.

**Table S4.** Differentially expressed genes between cells of wild-type and *Lmna^CKO^* hearts, derived from pseudo-bulk analysis of snRNA-seq data, related to **Figure 4**.

## METHODS

### Resource availability Lead contact

Kohta Ikegami.

### Materials availability

All mice are available at the sources specified in **Key Resources Table**. Plasmids produced in this study are provided upon request from the lead contact after material transfer agreements. Any information required to reanalyze the data reported in this paper is available from the lead contact.

### Data availability

High-throughput sequencing data associated with this version of the manuscript are limited to reviewers and journal editors. All data will become available to the public after peer review.

### Method details

#### Mouse genetics and treatment

*Lmna-LoxP* mice ^84^ were provided by Dr. Yixian Zheng at Carnegie Institution. *Myh6-MerCreMer* (Myh6MCM) transgene mice ^85^ (JAX stock No: 005657), *Rosa26^CAG-LSL-tdTomato^* (“Ai75”) mice ^86^ (JAX stock No: 25106), *Cgas-null* mice ^64^ (JAX stock No: 026554), and *CAG-Lox-Stop-Lox-Uprt* transgene mice ^87^ (*Uprt^Tg^*; JAX stock No: 021469) were obtained from the Jackson Laboratory. *Sting-null* mice were generated by breeding *Sting-flox* mice ^65^, obtained from the Jackson Laboratory (JAX stock No: 031670), with mice carrying the *Mef2C-AHF-Cre* transgene allele ^88^, provided by Dr. Brian L. Black at the University of California San Francisco. For cardiomyocyte-specific *Lmna* knockout experiments, *Lmna^F/F^;Myh6MCM^Tg/0^* (*Lmna^CKO^*) and *Lmna^+/+^;Myh6MCM^Tg/0^* (wild-type control) mice were used. For the analysis of nuclear tdTomato intensity, *Lmna^F/F^;Myh6MCM^Tg/0^*;*Rosa26^CAG-LSL-tdTomato/+^*and *Lmna^+/+^;Myh6MCM^Tg/0^*;*Rosa26^CAG-LSL-tdTomato/+^*mice were used. For SLAM-IT-seq experiments, *Lmna^F/F^;Myh6MCM^Tg/0^;Uprt^Tg/0^*, *Lmna^+/+^;Myh6MCM^Tg/0^;Uprt^Tg/0^*, *Lmna^F/F^;Myh6MCM^Tg/0^;Uprt^0/0^*, and *Lmna^+/+^;Myh6MCM^Tg/0^;Uprt^0/0^* mice were used. For *Cgas* deletion experiments, *Cgas^-/-^*;*Lmna^F/F^;Myh6MCM^Tg/0^* mice, *Cgas^+/+^*;*Lmna^F/F^;Myh6MCM^Tg/0^*, *Cgas^+/+^*;*Lmna^F/+^;Myh6MCM^Tg/0^*, and *Cgas^-/-^*;*Lmna^F/+^;Myh6MCM^Tg/0^* mice were used. For *Sting* deletion experiments, *Sting^-/-^*;*Lmna^F/F^;Myh6MCM^Tg/Tg^*mice, *Sting^+/+^*;*Lmna^F/F^;Myh6MCM^Tg/Tg^*, *Sting^+/+^*;*Lmna^F/+^;Myh6MCM^Tg/Tg^*, and *Sting^-/-^*; *Lmna^F/+^;Myh6MCM^Tg/Tg^* mice were used. All mice used in experiments were administered with tamoxifen (Sigma, T5648) at 6-8 weeks of age via intraperitoneal injections (100 μL of 4 mg/mL solution per day for 4 consecutive days) dissolved in corn oil (Sigma, C8267). All mice were used in mixed genetic backgrounds. All mouse experiments were approved by the Institutional Animal Care and Use Committee (IACUC) at Cincinnati Children’s Hospital under IACUC protocol 2021-0014 or at University of Chicago under IACUC protocol 71730-10. All procedures were performed in compliance with institutional and governmental regulations under PHS Animal Welfare Assurance number D16-00068 (Cincinnati Children’s) or D16-00322 (University of Chicago).

#### Mouse genotyping

For genotyping of the *Lmna* alleles, genomic DNA of isolated cardiomyocytes was extracted with Phenol/Chloroform/Isoamyl alcohol (Invitrogen,15593-031), treated with RNaseA and Proteinase K, and purified. Floxed, recombined, and wild type *Lmna* alleles were detected by PCR using the primers KI279 and KI280 (**Key Resources Table**). Based on the agarose gel (1.5%) analysis of PCR-amplified products, recombination efficiency (%) was calculated by 153-bp band (Recombined) over 598-bp band (Floxed) intensities. Genotyping of *Myh6-MerCreMer*, *Cgas*, *Sting*, *Uprt*, and *Rosa26::CAG-LSL-tdTomato* alleles was performed at Transnetyx, Inc.

#### Echocardiography

The transthoracic echocardiography was performed using a Vevo 3100 LT (FUJI FILM VisualSonics) and a transducer of 50-MHz MX-700. Parasternal long-axis view and short-axis view at the papillary muscle level were imaged. Two-dimensional M-mode tracing was recorded at three or more consecutive cardiac cycles. Data were analyzed using Vevo LAB Software Package V3.2.6. Echocardiographic parameters between two groups were compared using Wilcoxon rank sum tests, and P-values were adjusted for multiple testing using the Benjamini-Hochberg procedure in R.

#### Survival analysis

Kaplan-Meier survival analyses were performed using the *survival* package and visualized using the *survminer* package in R. Log-rank tests were performed using the *survdiff* function in the survival package.

#### Histological staining

Hearts were perfused with 100 mM KCl and fixed in 10% Formalin (Fisherbrand, 245-685). Fixed hearts were embedded in paraffin and sectioned to a thickness of 5 µm. Heart sections were stained with hematoxylin and eosin (H&E) or a Masson’s Trichrome Kit (Thomas Scientific LLC, KTMTRPT) according to the manufacturer’s protocol.

#### Cardiomyocyte isolation

Mice were administered with heparin (100 Units, NDC 25021-400-10), anesthetized with Isoflurane, and euthanized by cervical dislocation. Hearts were perfused with 100 mM KCl and moved to Tyrode’s solution (10 mM glucose, 5 mM HEPES, 5.4 mM KCl, 1.2 mM MgCl_2_, 150 mM NaCl and 2 mM sodium pyruvate, pH 7.35). Excised hearts were cannulated to the Langendorff retrograde perfusion system through the aorta and perfused with Base solution (Tyrode’s solution with 10 mM taurine and 12 mM 2,3-butanedione monoxime) and then digested with prewarmed Digestion buffer (Base solution with 180 U/mL Collagenase Type II (Worthington, LS004177) and 25 μM CaCl_2_) for 20 minutes at 37°C. Heart tissue was then isolated, gently minced in Base solution containing 5 mg/mL BSA. Cell suspension was filtrated through a 240 μm mesh. Cardiomyocytes settled to the bottom of conical tubes by gravity were used in experiments.

#### Immunohistochemistry

Deparaffinized heart tissue sections, that had been fixed with 10% formalin, were subjected to heat-induced antigen retrieval in Target Retrieval Solution (S2367, DAKO) for 5 minutes. Antigen-retrieved sections were incubated with anti-Lamin A/C antibody Ab26300 (rabbit, Abcam) (1:400) for 1 hour at room temperature, followed by biotinylated anti-rabbit IgG antibody for 30 minutes at room temperature. The antigen-antibody binding was detected by Elite kit (PK-6100, Vector Laboratories) and DAB (DAKO, K3468) system. Sections were counterstained by hematoxylin.

#### Immunofluorescence on tissue sections

Deparaffinized heart tissue sections, that had been fixed with 10% formalin, were subjected to heat-induced antigen retrieval in 10 mM Tris EDTA buffer (pH 9.0) for 5 minutes. Antigen-retrieved sections were incubated overnight at 4 °C with goat anti-CD45 antibody (R&D Systems, AF114-SP) and rabbit anti-CD68 antibody (Cell Signaling, 97778), or rabbit anti-γ-H2A.X antibody (Cell Signaling, 9718) and goat anti-desmin antibody (Invitrogen, PA5-19063), or rabbit anti-PCM1 antibody (Sigma, HPA023370), goat anti-TdTomato antibody (OriGene, AB8181), and mouse anti-Cox4 antibody (R&D Systems, MAB6980), followed by Alexa fluorophore-conjugated secondary antibodies for 1 hour at 37°C. Cells were counterstained with DAPI (4′,6-diamidino-2-phenylindole), submerged in 0.25% Sudan Black B (Electron Microscopy Slides, #21610) in 70% Isopropanol for 10 minutes, and then mounted with ProLong Glass Antifade Mountant (Invitrogen, P36984). Fluorescence signals and imaging were acquired using a Nikon A1R laser-scanning confocal microscope. CD45 and CD68 quantification of whole heart tissue was performed using Nikon NIS-Elements; each genotype had five biological replicates. CD45/CD68 positive cells were identified based on the overlap of the DAPI signal with CD45/CD68. Signal intensities between two groups were compared using Wilcoxon rank sum tests in R. For γ-H2A.X quantification, nuclei with DAPI were counted and those with at least one γ-H2A.X focus were quantified using CellProfiler. Five to eight images per mouse (374 x 374 µm^2^ per image) were used for the quantification.

#### Immunofluorescence on isolated cardiomyocytes

Isolated cardiomyocytes were fixed in 4% paraformaldehyde (Electron Microscopy Slides, #15710) in PHEM buffer (60 mM PIPES pH7.5, 25 mM HEPES pH7.5, 10 mM EGTA, 4 mM MgSO_4_) for 10 minutes at 37°C, then washed with PBS and attached to coverslips with Cell-Tak Cell Adhesive (Sigma Aldrich, 354240). Cells were blocked and permeabilized with in a buffer containing 5% normal donkey serum (Jackson ImmunoResearch, 017-000-121), 1% non-fat milk, and 0.1% Triton X-100 in PBS for 1 hour at 37°C. Permeabilized cells were incubated overnight at 4°C with mouse anti-Lamin A/C antibody (Santa Cruz, sc-376248), rabbit anti-PCM-1 antibody (Sigma, HPA023370), mouse anti-cGAS antibody (Cell Signaling, D3O8O). Cells were washed and incubated with Alexa fluorophore-conjugated secondary antibodies and Alexa Fluor Plus 647 Phalloidin for 1 hour at 37°C. Cells were counterstained with DAPI and then mounted with ProLong Glass Antifade Mountant. Fluorescence signals were detected on Nikon A1R laser-scanning confocal microscope or Yokogawa CSU-W1 Sora spinning disk confocal microscope.

#### Plasmid construction

The expression vectors used in this study were derived from the pAAV:cTNT::Luciferase vector (gift from William Pu; Addgene plasmid # 69915) ^89^. For cloning of pAAV:cTnT::cGAS and pAAV:cTnT::GFP vectors, we digested the pAAV:cTNT::Luciferase vector with *NheI* and *NotI*, and the luciferase gene was replaced with the cGAS or EGFP sequences using NEBuilder HiFi DNA Assembly Master Mix (NEB E2621). The pAAV:cTnT::GFP-icGAS vector was generated by inserting the catalytically-inactive cGAS (icGAS) sequence (DNA fragment ID: EA086, **Key Resources Table**) into the pAAV:cTnT::GFP vector digested with *NotI*. The icGAS sequence was derived from the human cGAS protein sequence with an E225A/D227A amino acid substitution that abolishes enzyme activity and interferon production, but retains DNA binding ability and functions, used as a NE rupture marker previously ^57^. The DNA fragments for mouse wild-type *Cgas* (NM_173386.5) and *icGAS* were synthesized using gBlocks Gene Fragment synthesis service (Integrated DNA Technologies). *EGFP* with a flexible GS linker (GGGGS) at the C-terminus ^90^ was amplified from the pLJM1-EGFP vector (gift from David Sabatini; Addgene plasmid #19319) ^91^ using the EA062 and EA064 primers (**Key Resources Table**).

#### MyoAAV production and *in vivo* transduction

For myoAAV production, we used a published protocol with modifications ^92^. We co-transfected the pAAV-cTnT-transgene vector (see above), the pHelper vector (GenBank: AF369965.1), and the pRep/Cap 1A-MYO capsid vector ^56^ into AAVpro 293T cells (TaKaRa, 632273) using polyethylenimine (PEI) reagent. The 1A-MYO capsid vector allows production of muscle-tropic AAV9 derivative (MyoAAV)^56^. The transfected cells were cultured in the OptiPRO^TM^-SFM medium (Thermo Fisher, 12309019) for 72 hours at 37°C to produce virus. Cells and the culture media were collected for AAV purification. AAV particles were extracted with chloroform, precipitated with polyethylene glycol, purified with DNase/RNase digestion followed by chloroform extraction, and concentrated in PBS with Amicon filters (Millipore), as detailed in a previous report ^92^. The viral genome copy numbers (vg) were estimated by qPCR using primers amplifying the cTnT promoter region (primer ID KI466 and KI467, **Key Resources Table**). MyoAAV particles were administered to mice through retro-orbital venous sinus injection at a dose of 2×10^11^ vg per mouse at 4 weeks of age, which was 14-15 days prior to tamoxifen.

#### Image quantification related to nuclear envelope ruptures

For quantification of nuclei with PCM1-lost nuclear tips and DNA protrusion, we used immunofluorescent images of isolated cardiomyocytes stained for PCM1 and counter-stained with DAPI (DNA) and Phalloidin (F-actin). Rod-shape cardiomyocytes that had a well-defined sarcomere structure based on Phalloidin staining and two or more nuclei were used for quantification. We quantified instances of DAPI signal protrusion from a nuclear envelope site at which PCM1 signals were specifically discontinued (“local PCM1 loss”). The percentage of cells with one or more nuclei with a local PCM1 loss and the number of such nuclei per cell were quantified manually post imaging. We obtained the percentage from >50 cardiomyocytes per animal with a total of 3 animals per genotype. The percentages of cardiomyocytes with PCM1-lost nuclei were compared between genotypes using unpaired one-tailed Welch’s t-tests in R.

For quantification of nuclei with icGAS puncta, we used isolated cardiomyocytes from wild-type or *Lmna^CKO^* mice treated with MyoAAV-GFP-icGAS. The percentage of cells with icGAS puncta was quantified manually post imaging. We obtained the percentage from >50 cardiomyocytes per animal with a total of 3 animals per genotype. The icGAS-positive cardiomyocyte percentages were compared between genotypes using an unpaired one-tailed Welch’s t-test in R.

For quantification of NLS-tdTomato intensities, we used *Lmna^F/F^;Myh6MCM^Tg/0^*;*Rosa26^CAG-LSL-tdTomato/+^* and *Lmna^+/+^;Myh6MCM^Tg/0^*;*Rosa26^CAG-LSL-tdTomato/+^*mice. Isolated cardiomyocytes were fixed, immunostained for PCM1, and counterstained with DAPI. Quantification of NLS-tdTomato signals was performed in the Fiji image analysis software. Cardiomyocytes that had two or more nuclei were used for quantification. Nuclei with nuclear envelope ruptures were identified as described above. A freehand-drawn area was placed inside a nucleus for quantification of the nuclear tdTomato signal. A nucleus was selected for quantification using the following algorithm: If a cell had only ruptured nuclei, one ruptured nucleus was randomly selected for quantification; if a cell had both ruptured and unruptured nuclei, one ruptured nucleus was randomly selected for quantification; if a cell had only unruptured nuclei, one unruptured nucleus was randomly selected for quantification. For each nucleus, whether it had a local PCM1 loss with DNA protrusion was manually determined as described above. A second quantification area of the same size as the nuclear area was placed in the cytoplasm for cytoplasmic tdTomato signal quantification. For each of the two areas, a mean NLS-tdTomato pixel intensity was measured using the ROI manager function in Fiji. We performed this quantification process for three mice per genotype. In total, we had quantification information for 99 wild-type cardiomyocytes (all intact PCM1), 90 *Lmna^CKO^* cardiomyocytes with local PCM1 loss, and 36 *Lmna^CKO^* cardiomyocytes with intact PCM1. Signal intensities between two groups were compared using Wilcoxon rank sum tests, and P-values were adjusted for multiple testing using the Benjamini-Hochberg procedure in R.

#### TUNEL assay

TUNEL assays were performed on deparaffinized heart sections using a Click-iT™ Plus TUNEL Assay Kits for In Situ Apoptosis Detection (Invitrogen, C10617), according to the manufacturer’s instruction. The samples were then co-stained with Alexa Fluor Plus 647 Phalloidin (Invitrogen, A30107) for 1 hour at 37°C to visualize cells in tissue sections. For positive controls, sample tissues were treated with DNase I prior to the TUNEL assay. The number of cells with positive staining in the TUNEL assay (TUNEL^+^ cells) was determined using sections from three hearts per genotype. TUNEL positive cells in 2-3 images per mouse (257 x 257 µm^2^ per image) were counted manually and compared between two groups using Wilcoxon rank sum tests in R.

#### Immunoblot

Snap-frozen hearts or isolated cardiomyocytes were lysed in the RIPA buffer for cGAS and STING immunoblotting (20 mM Tris-HCl, 150 mM NaCl, 1% IGEPAL, 0.5% sodium deoxycholate, 0.1% SDS, 10 mM dithiothreitol, protease and phosphatase inhibitors) or the Urea buffer for Lamin A/C immunoblotting (20 mM HEPES pH 7.4, 1 M NaCl, 8 M urea, protease and phosphatase inhibitors). Proteins were extracted using pestle homogenization and sonication. Cell lysates were centrifuged at 13,000 rpm for 5 minutes at 4°C, and proteins in the supernatant were separated by SDS-PAGE. Proteins were transferred to a PVDF membrane. Membranes were blocked with nonfat milk. The primary antibodies were rabbit anti-Lamin A/C antibody (Santa Cruz, sc-20681, 1:1000), rabbit anti-cGAS antibody (Cell Signaling, #31659, 1:500), rabbit anti-STING antibody (Cell Signaling, #13647, 1:500), and rabbit anti-GAPDH antibody (ABclonal, AC001, 1:1000); Secondary antibodies were anti-rabbit or anti-mouse IgG (H+L) DyLight 680 and 800 (Cell Signaling, 1:5000). Signals were detected and quantified in the Odyssey CLx Imager (LI-COR). Gels after protein transfer were counter-stained with Coomassie to evaluate the loaded protein amount.

#### Electron microscopy

Heart tissues were dissected and fixed in 2.5% glutaraldehyde, post-fixed in 1% OsO4 for 1 hour at 4°C, rinsed, dehydrated in ethanol, and infiltrated overnight. The semithin Epon sections were screened by light microscopy to select areas with longitudinal orientation of myocytes that were subsequently processed to prepare thin sections. The sections were stained with 1% uranyl acetate, followed by lead citrate. The sections were imaged using Hitachi Model H-7650 Transmission Electron Microscope at Cincinnati Children’s Integrated Pathology Research Facility.

#### Quantitative reverse-transcriptase PCR (RT-PCR)

For RT-PCR analysis, we used wild-type treated with MyoAAV-Luciferase (n=5) or MyoAAV-cGAS (n=3) and *Lmna^CKO^* mice treated with MyoAAV-Luciferase (n=4) or MyoAAV-cGAS (n=5) at 2 weeks post tamoxifen. Total RNAs of isolated cardiomyocytes were extracted with Trizol LS (Invitrogen, 10296010), treated with DNaseI, and purified. Quantitative RT-PCR was conducted with the Luna Universal One-Step RT-qPCR Kit (NEB, E3005) on a Bio-Rad CFX96 Real-Time PCR Detection System (Bio-Rad) in accordance with the manufacturer’s instructions. The primers for mouse genes were: EA096 and EA097 for *Cxcl10*; EA098 and EA099 for *Ifnb1*; EA100 and EA101 for *Ifit3*; and KI444 and KI445 for *Actb* (**Key Resources Table**). The delta-delta Ct method was used to quantify mRNA abundance with *Actb* as a normalization control. mRNA levels between two groups were compared using Wilcoxon rank sum tests, and P-values were adjusted for multiple testing using the Benjamini-Hochberg procedure in R.

#### RNA-seq

For RNA-seq comparing wild-type mice with *Lmna^CKO^* mice at different post tamoxifen times, we used wild-type (n=5) and *Lmna^CKO^*mice (n=4) at 1 week post tamoxifen, wild-type (n=3) and *Lmna^CKO^*mice (n=2) at 11 days post tamoxifen, and wild-type (n=7) and *Lmna^CKO^* mice (n=5) at 2 weeks post tamoxifen. For RNA-seq analysis of *Lmna/Cgas* double knockout mice, we used *Cgas^-/-^*;*Lmna^F/F;Myh6MCM^* mice (n=4), *Cgas^+/+^*;*Lmna^F/F;Myh6MCM^* (n=6), *Cgas^+/+^*;*Lmna^F/+;Myh6MCM^* (n=6), and *Cgas^-/-^*;*Lmna^F/+;Myh6MCM^*(n=4) for experiments. Hearts were perfused with 100 mM KCl and removed for RNA extraction. Total RNAs were extracted with Trizol LS (Invitrogen, 10296010), treated with DNaseI, and purified. For tamoxifen time course experiments, mRNA sequencing libraries were generated using the poly-A selection module in the NEBNext UltraII Directional RNA Library Prep Kit (NEB E7760) and sequenced on the Illumina HiSeq 2500 sequencer with single-end 50 cycles. For *cGas*/*Lmna* double knockout experiments, libraries were generated using QuantSeq 3’ mRNA-Seq Library Prep Kit FWD (Lexogen, K01596) and sequenced on the Illumina NovaSeq 6000 sequencer with single-end 100 cycles.

#### RNA-seq analysis

High-throughput sequencing reads were aligned to the mouse mm39 reference genome with the Gencode vM27 basic gene annotation using STAR version 2.7.9 ^93^ with the default alignment parameters except using “clip3pAdapterSeq AGATCGGAAGAGCACACGTCTGAACTCCAGTCA”. From raw read counts per transcript, TPMs (transcripts per million) were calculated in R as follows: TPM=10^6^ x RPK/sum(RPK), where RPK=read count/transcript length in kp, and sum(RPK) is the sum of RPKs for all transcripts. Raw read counts were used for differential gene expression analyses using DESeq2 ^94^ in R. Protein-coding or lncRNA genes with adjusted P-value < 0.05 and absolute log_2_ fold change > 1 (NEB Directional RNA-seq) or with adjusted P-value < 0.05 and absolute log_2_ fold change > 0 (QuantSeq 3’ mRNA-seq) were considered differentially expressed genes (**Table S1**). Differentially expressed genes were analyzed for the enrichment of GO Biological Processes, GO Molecular Functions, GO Cellular Components, and KEGG pathway terms using Metascape ^95^ with the default enrichment parameters.

#### Single-nucleus RNA-seq

For single-nucleus RNA-seq, we used wild-type (n=3) and *Lmna^CKO^*mice (n=3) at 2 weeks post tamoxifen. Combinatorial indexing-based single-nucleus RNA-seq was conducted according to the optimized sci-RNA-seq3 protocol developed previously ^96^. Briefly, mice were anesthetized with isoflurane, euthanized by cervical dislocation, and then hearts were perfused with 100 mM KCl. Removed hearts were diced into small pieces in cold PBS containing Diethyl pyrocarbonate (DEPC, Sigma D5758). Fresh heart pieces were dounce-homogenized to liberate nuclei in Hypotonic lysis buffer B (7.7 mM Na_2_HPO_4_, 4.5 mM NaH_2_PO_4_, 1.8 mM KH_2_PO_4_, 2.7 mM KCl, 10.3 mM NaCl, 3 mM MgCl_2_, 0.08% BSA, 0.025% Igepal CA-630, 1% DEPC) and passed through cell strainers. Nuclei were fixed in 1 mg/mL dithiobis(succinimidyl propionate) (DSP, Thermo Fisher 22585) for 15 min on ice, washed, and stored at –80°C until use. For each mouse, 50,000 nuclei were placed in each of 8 wells of a 96-well plate (total 48 wells for 6 mice). The remaining 48 wells were filled with similarly processed mouse heart nuclei with comparable sample quality from an unrelated project. Nuclei were subjected to reverse transcription with 96-indexed poly-T RT primers that contained a unique molecular identifier (UMI) sequence (3lvl_mRNA_RT_plate_1; **Table S3**). Nuclei from all wells were combined and redistributed evenly to 96 wells. Redistributed cells were subjected to ligation with 96-indexed ligation primers in 96 wells (3lvl_mRNA_Lig_plate_1; **Table S3**). Nuclei from all wells were combined again and redistributed evenly to 96 wells at 1,000 nuclei per well. Redistributed cells underwent second strand synthesis, protease digestion, tagmentation using N7-loaded Tn5 transposase, and PCR-amplified using 96-indexed PCR P7 primers (PCR_P7_plate1; **Table S3**) and the unindexed P5 primer, for 16 cycles. PCR amplicons were combined and size-selected for DNA fragments between 250 bp and 600 bp. Purified library DNA were sequenced for 50 bases on the paired-end mode (34 cycles on Read1 to sequence the ligation index, UMI, and the RT barcode; 10 cycles on Index1 to sequence the P7 index; and 48 cycles on Read2 to sequence the cDNA) using an Illumina NovaSeq6000 sequencer.

#### Single-nucleus RNA-seq analysis

Pre-processing of the single-nucleus RNA-seq data was performed using the published sci-RNA-seq3 pipeline ^96^, available in Github (https://github.com/JunyueC/sci-RNA-seq3_pipeline). The sci-RNA-seq3 pipeline takes fastq files separated by the P7 index as input, aligns reads to the genome, and returns a gene-by-cell read count matrix. First, raw reads were trimmed to remove poly A tails, aligned to the mm39 mouse reference genome using STAR, and filtered for MAPQ greater than or equal to 30. Duplicate reads were removed based on UMI. Read counts per gene were computed for each cell defined by the unique combination of the RT index, ligation index, and P7 index. The gene-by-cell count matrix produced by the sci-RNA-seq3 pipeline was further processed in the *SingleCellExperiment* package framework in R^97^. We used the *scDblFinder* package for doublet detection and the *scuttle* package for mitochondrial RNA quantification in R. We obtained 14,111 nuclei after filtering for nuclei with the *scDblFinder* doublet score less than 0.25 and mitochondrial RNA contamination percentage less than 2% of total RNAs per cell. Read counts for a gene were normalized by per-cell read depth and then log-transformed to obtain normalized expression values (“logcounts”). Logcounts were used to plot expression values in graphs. Principal component analysis (PCA) was performed on logcounts for 2,000 highly variable genes across cells using the *fixedPCA* function in the *scran* package. The top 50 principal components were used in the *runUMAP* function in the *scater* package with n_neighbors=20 and min_dist=1 parameters to obtain UMAP dimensions. To cluster cells, logcounts were processed in the *clusterCells* function in the *scran* package using the top 50 principal components, with the number of nearest neighbors k=12. Cell clusters were annotated manually with cell type names based on marker gene expression. Per-gene read counts were summed within cell groups and processed by the *DESeq2* package ^94^ to perform pseudo-bulk differential gene expression analysis. Genes with adjusted P-values smaller than 0.05 were defined as differentially expressed genes (**Table S4**). GO analysis of differentially expressed genes was performed as described in the RNA-seq analysis section. For the analysis for the enrichment within selected GO and KEGG terms, all genes associated with the selected terms were obtained using the R packages *GO.db* and *KEGGREST*, and enrichment of differentially expressed genes among these genes was computed by Fisher’s exact test with P-values adjusted by the Benjamini-Hochberg procedure. Cell-cell communication analysis was performed using the *CellChat* package in R ^74^ using the 12 cell type annotations and the default mouse database with a minimum of ten cells for the analysis. Positive communications were selected by the *subsetCommunication* function with threshold P-value less than 0.05, and the reported interaction counts were used to plot data.

#### SLAM-IT-seq

For SLAM-IT-seq experiments, we used *Lmna^F/F^;Myh6MCM^Tg/0^;Uprt^Tg/0^*(n=6) and *Lmna^+/+^;Myh6MCM^Tg/0^;Uprt^Tg/0^* (n=8) mice as experimental groups, and *Lmna^F/F^;Myh6MCM^Tg/0^;Uprt^0/0^*(n=4) and *Lmna^+/+^;Myh6MCM^Tg/0^;Uprt^0/0^* (n=3) mice as negative control groups that would not incorporate 4-thiouracil (4sU) due to the absence of the *Uprt* allele. All mice were administered with tamoxifen and used at 2 weeks post tamoxifen treatment. SLAM-IT-seq was conducted according to a previously established protocol ^53^. Briefly, mice were intraperitoneally injected with 4-thiouracil (4sU) dissolved in a DMSO/corn oil (1:3) solution at a dose of 10 mg per gram of body weight. Six hours after the injection, mice were sacrificed, and hearts were perfused with 100 mM KCl and removed. RNAs from the left ventricular free wall were extracted by Trizol LS, purified, and DNaseI-treated using Directzol RNA miniprep kit (Zymo, R2052). Purified RNAs (1.5 µg) were subjected to alkylation in the presence of 12 mM Iodoacetamide (IAA, Sigma I1149). High-throughput sequencing libraries were generated from the alkylated RNAs using Quant-seq 3’ mRNA-seq FWD kit (Lexogen K01596) and sequenced 100 bases on the single-end mode on an Illumina NovaSeq6000 sequencer.

#### SLAM-IT-seq analysis

We analyzed SLAM-IT-seq data according to the published SLAM-IT-seq data analysis pipeline that uses the *slamdunk* package ^98^, available on GitHub (https://github.com/t-neumann/slamdunk), and R functions^59^. First, raw reads were processed by the *slamdunk all* function in the the *slamdunk* package with the following alignment parameters: --trim-5p 12 --topn 100 --multimap --max-read-length 101 with default -- var-fraction (0.8) and default -c (1). This function aligned reads to 3’ UTRs of Gencode vM27 genes (total 208,307 UTRs from 37,065 unique genes) in the mm39 mouse reference genome. Aligned reads were processed by the *alleyoop utrrates* function in the *slamdunk* package with default parameters to quantify C-to-T conversion events. UTRs with at least one read coverage for all 21 samples were retained for further analyses (47,717 UTRs from 14,250 unique genes). Statistically significant C-to-T conversion events in each transcript (i.e. the likelihood of cardiomyocyte-derived transcripts) were identified by comparing the conversion events in mice carrying the *Uprt* allele with those in mice not carrying this allele using beta-binomial test (*bbtest* function) in R. P-values of the beta binomial test were adjusted with the Benjamini-Hochberg procedure for multiple tests (**Table S2**). A transcript with the lowest adjusted P-value was chosen to represent a gene if there were multiple transcripts per gene. Genes with the adjusted P-values smaller than 0.05 were defined as originating from cardiomyocytes, whereas genes with adjusted P-values greater than or equal to 0.05 were defined as not originating from cardiomyocytes.

#### Quantification and statistical analysis

Statistical details of experiments and sample sizes are described in Method sections and indicated in figures and/or figure legends. Quantitative data were shown as means (bar) of biological replicates (dots). P values < 0.05 were considered statistically significant. P-values are indicated in the figures as appropriate.

#### Image graphics

Graphical abstract and the study summary (**Fig.4J**) were created with BioRender.com.

## Key resources table

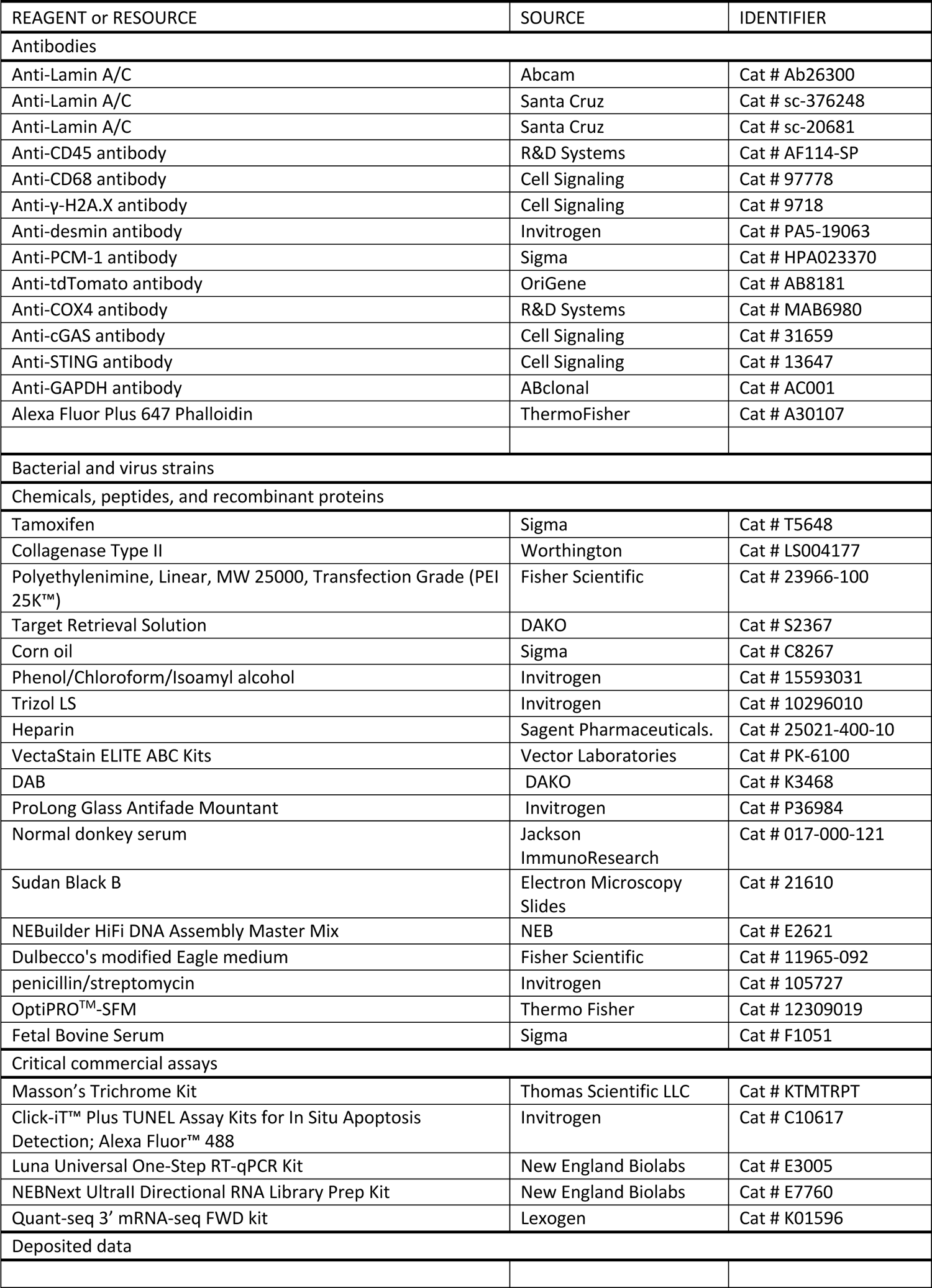
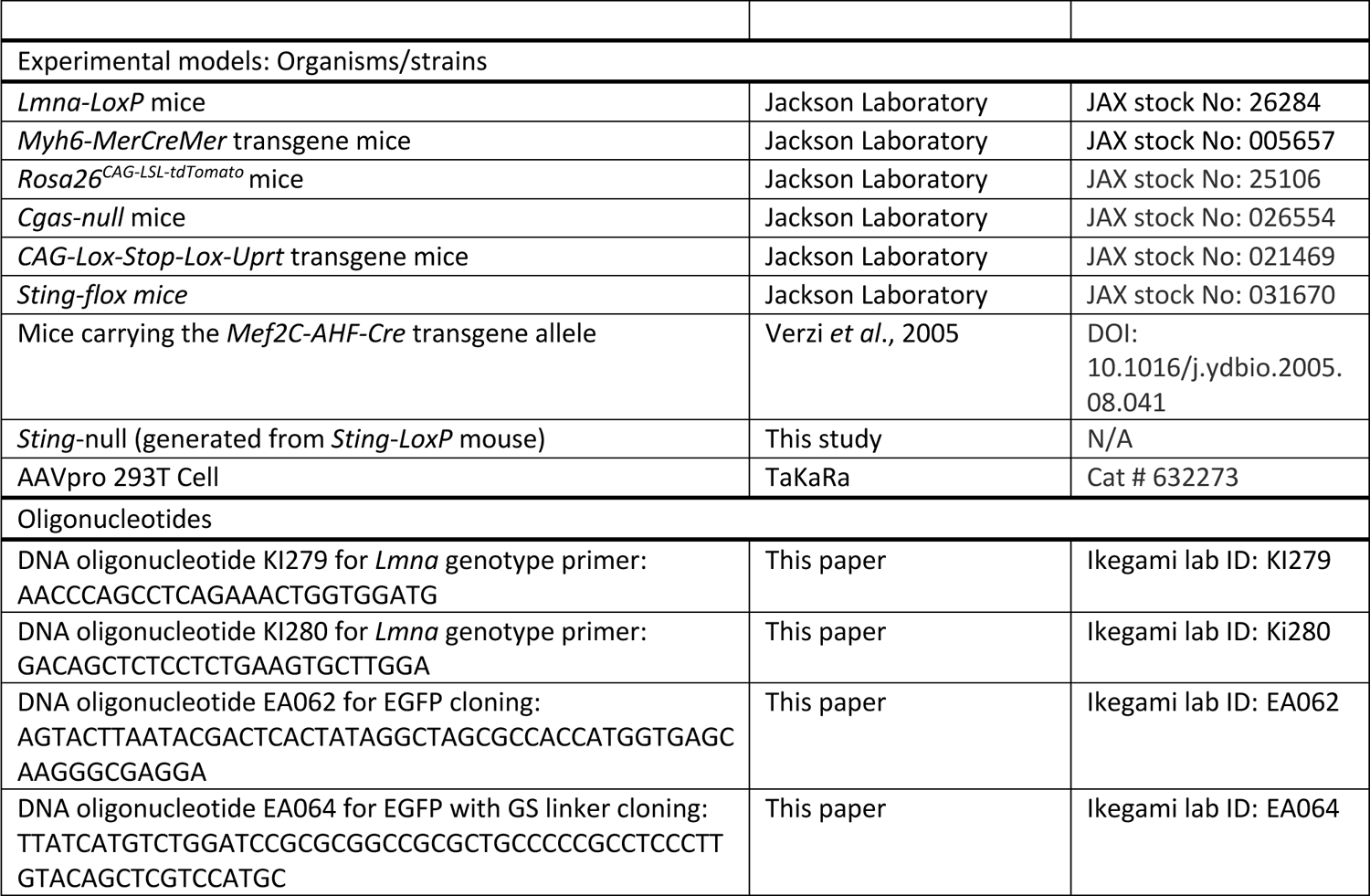
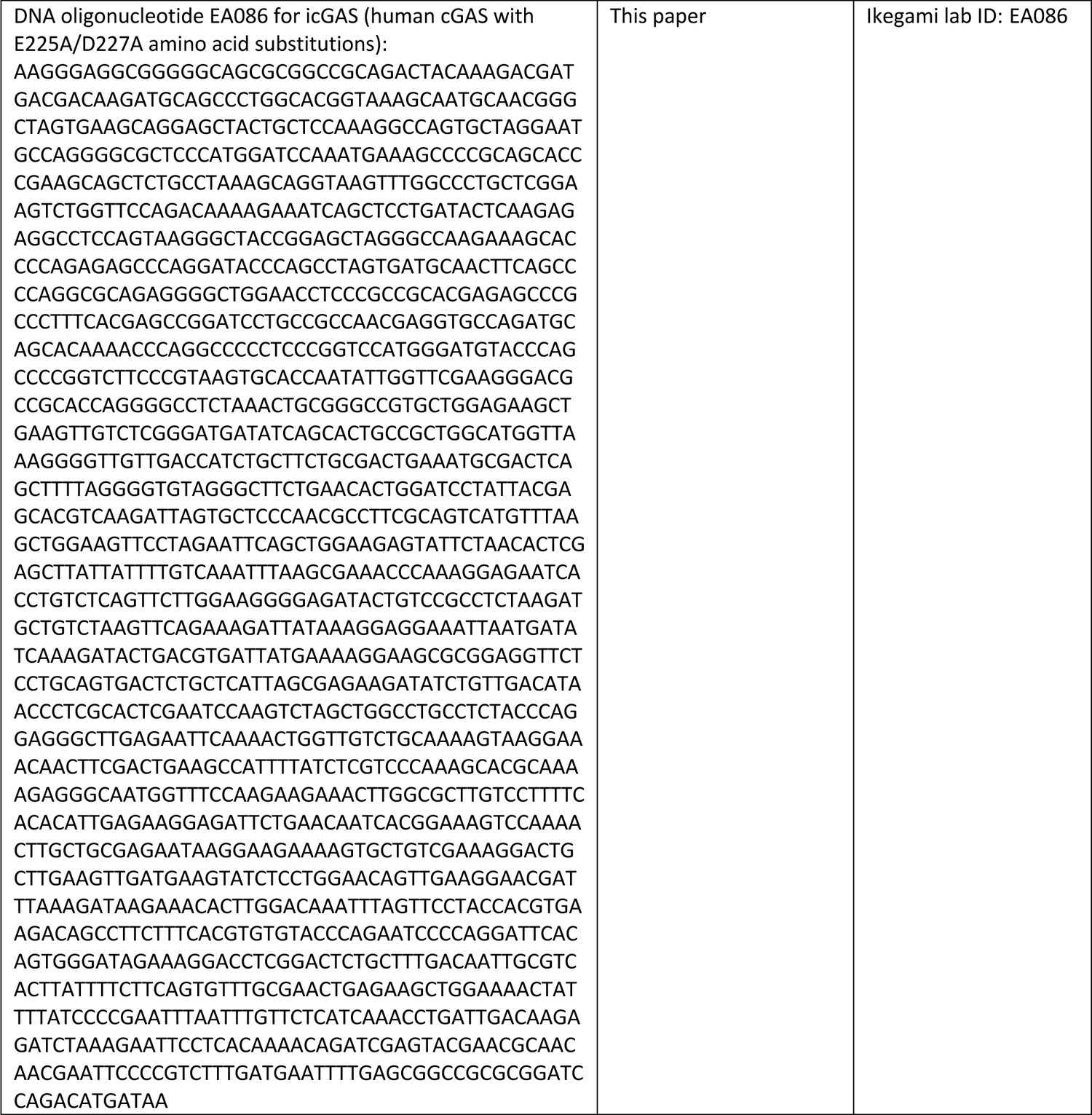
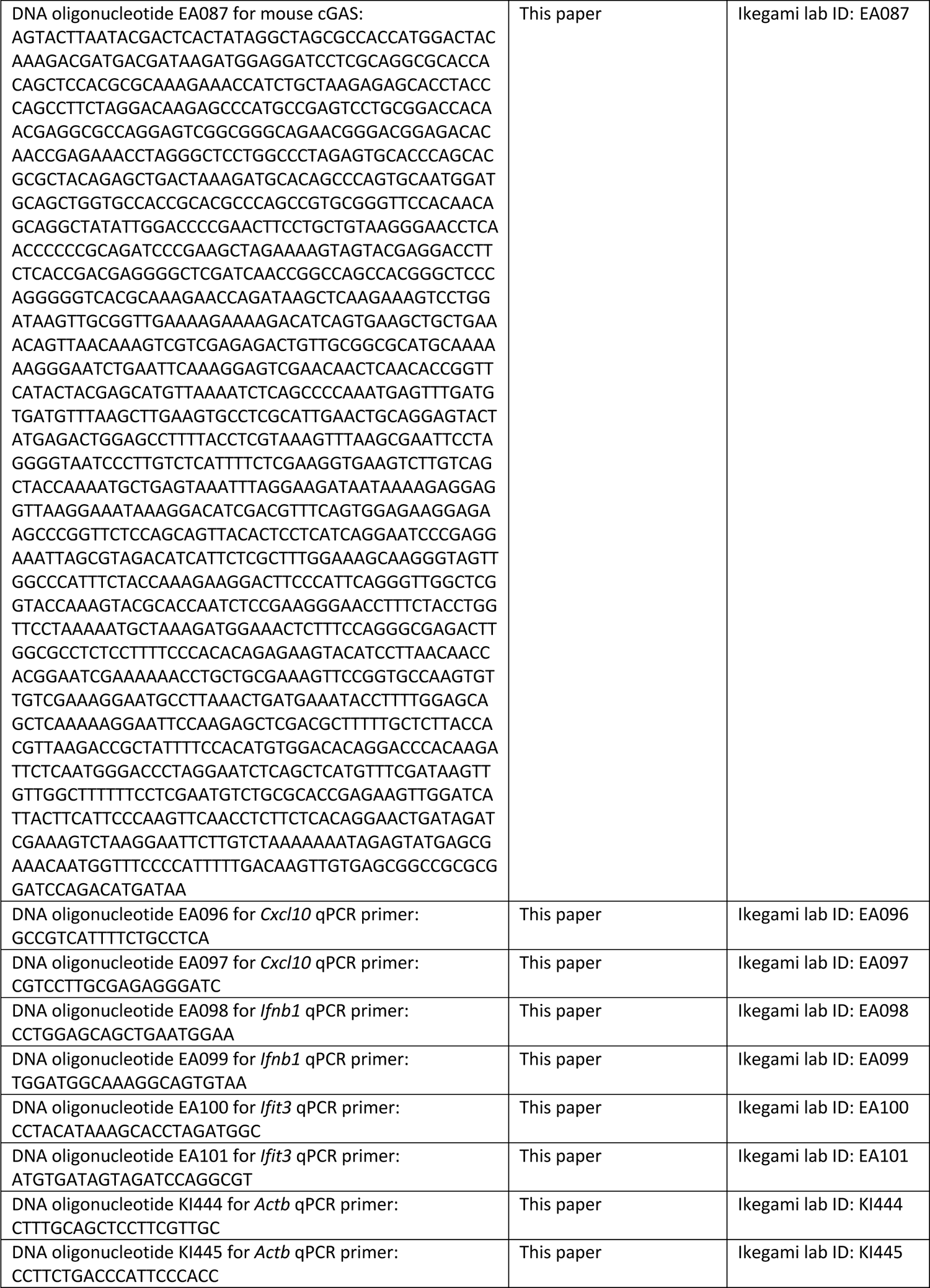
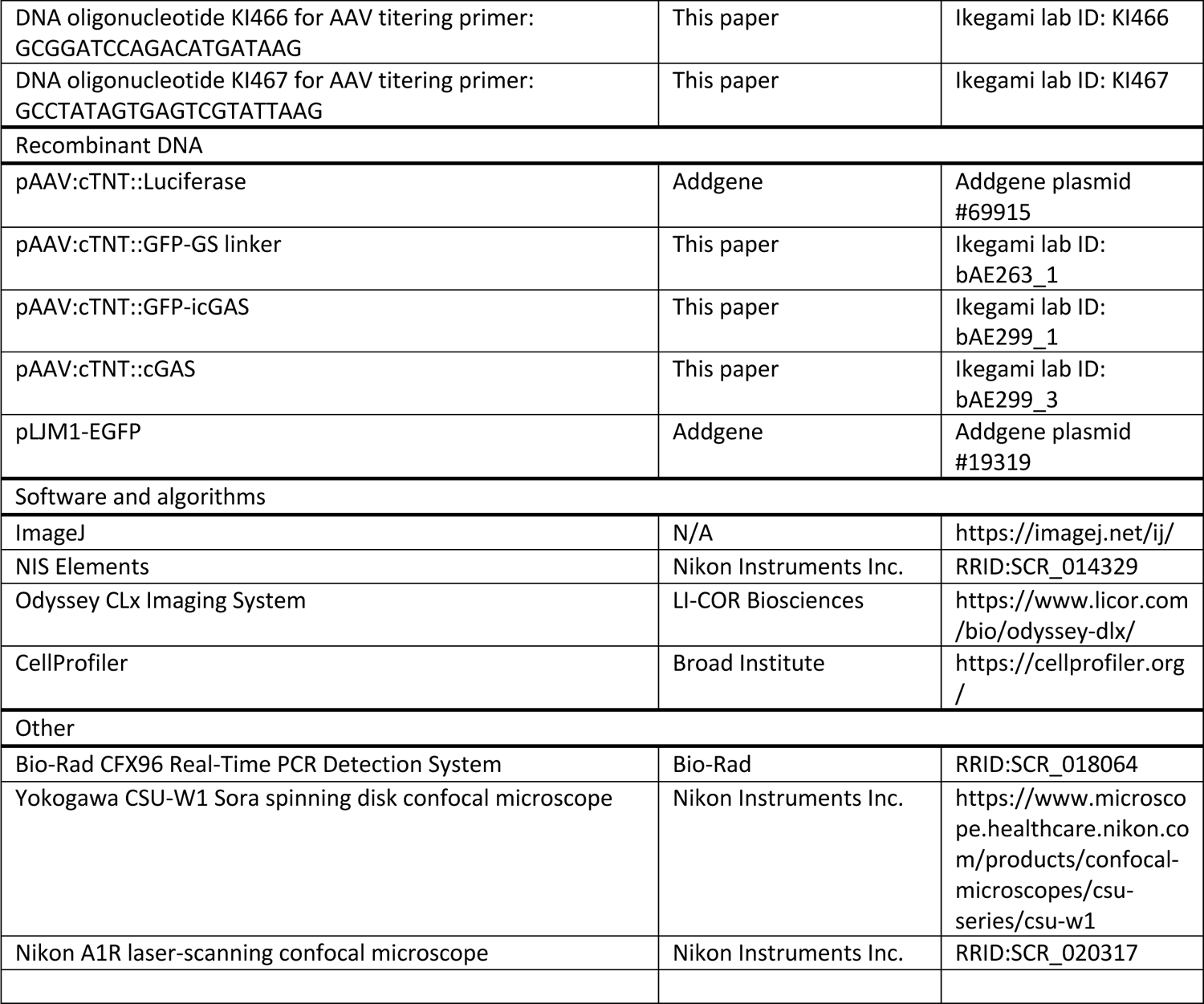

